# Acute lymph node slices are a functional model system to study immunity ex vivo

**DOI:** 10.1101/865543

**Authors:** Maura C. Belanger, Alexander G. Ball, Megan A. Catterton, Andrew W.L. Kinman, Parastoo Anbaei, Benjamin D. Groff, Stephanie J. Melchor, John R. Lukens, Ashley E. Ross, Rebecca R. Pompano

## Abstract

The lymph node is a highly organized and dynamic structure that is critical for facilitating the intercellular interactions that constitute adaptive immunity. Most ex vivo studies of the lymph node begin by reducing it to a cell suspension, thus losing the spatial organization, or fixing it, thus losing the ability to make repeated measurements. Live murine lymph node tissue slices offer the potential to retain spatial complexity and dynamic accessibility, but their viability, level of immune activation, and retention of antigen-specific functions have not been validated. Here we systematically characterized live murine lymph node slices as a platform to study immunity. Live lymph node slices maintained the expected spatial organization and cell populations while reflecting the 3D spatial complexity of the organ. Slices collected under optimized conditions were comparable to cell suspensions in terms of both 24-hr viability and inflammation. Slices responded to T cell receptor cross-linking with increased surface marker expression and cytokine secretion, in some cases more strongly than matched lymphocyte cultures. Furthermore, slices processed protein antigens, and slices from vaccinated animals responded to ex vivo challenge with antigen-specific cytokine secretion. In summary, lymph node slices provide a versatile platform to investigate immune functions in spatially organized tissue, enabling well-defined stimulation, time-course analysis, and parallel read-outs.

---

Events in the lymph node (LN) determine whether a host successfully fights infection and responds to vaccines, whether a nascent tumor is recognized and destroyed, and whether host tissues remain safe from autoimmunity. These immune responses arise in large part from precise spatial organization of cells and proteins in the lymph node.^1–3^ The structure of a lymph node can be roughly divided into an outer cortex containing B cell follicles, a T cell zone, or paracortex, and an inner medulla.^1,4^ During an immune response, cells in these regions communicate through both physical contact and secreted signals. Diffusion and the formation of gradients of secreted cytokines through the extracellular matrix generate orchestrated cell migration,^5,6^ and the local concentration of cytokines and other signals can drive strong positive feedback and divergent outcomes effecting the overall health of the host.^7^ All of these features suggest that the organization of the node may be essential to its function,^2,8^ and indeed, many similarly complex non-linear biological systems are exquisitely sensitive to spatial organization.^9^

Investigating the function of the lymph node with high spatial, temporal, and chemical resolution within a realistic microenvironment is challenging with existing experimental systems. Recent technological advances in immunological analysis have focused significantly on high-content single cell data using flow cytometry or mass cytometry,^10,11^ analysis of single-cell secretion and gene expression using microfluidics,^12–15^ and on bulk measurements such as metabolomics^16^ and live cell metabolic analysis.^17^ However, these cannot provide information on LN organization. Complementing this work, live in vivo imaging was developed over 15 years ago and continues to provide impressive insight into the dynamics of cell and tissue-level behavior in the native environment.^18–22^ Yet, it is challenging to experimentally manipulate tissues in vivo without prior genetic modification (e.g. optogenetics) or invasive injection. Approaches that retain the tissue’s spatial organization via fixation have revealed distinct regional subpopulations of cells,^23–25^ but fixed tissue is not amenable to experimental manipulation. While existing technologies can reveal important aspects of LN biology, a single approach that maintains the biological complexity of the organ while providing dynamic experimental access close to that of traditional cell cultures is still largely missing from the immunologist’s toolbox.^26^

Live ex vivo slices of lymph node tissue may provide a necessary middle-out approach, in a manner complementary to in vitro and in vivo work. Decades of work with brain slices^27,28^ set a precedent for both acute^29,30^ and long-term experimentation^31^, which informed protocols for other tissues such as the pancreas,^32^ liver,^33^ lung,^34^ and heart^35^. Unlike in vitro cell culture, slices cultured ex vivo preserve the extracellular microenvironment and any stromal and matrix-bound signals, which are essential to proper cellular positioning and motility.^2,5,36–38^ Furthermore, all cell types are retained in their correct ratios, whereas standard tissue dissociation (crushing and filtering) selectively depletes matrix-bound populations such as dendritic cells.^39^ In contrast to in vivo work, using tissue slices simplifies timecourse analysis via repeated measurements of the same tissue sample, especially after ex vivo stimulation.^40–44^ Slices also allow for precise stimulation of the organ interior, because precise quantities of drugs or other agents can be added at known concentrations and at known times.^45,46^ Furthermore, slices can be coupled together or co-cultured to generate a simplified model of inter-organ communication, akin to multi-organ organ-on-chip systems used to model pharmacokinetics and disease mechanisms.^47,48^ In the field of immunology, slices of the thymus have been used extensively to study T cell development.^21,45,49^ Live spleen^50,51^ and tonsillar^40,52,53^ slices were demonstrated 20 years ago, and continue to be a valuable tool to study immune function and viral infection. Live slices of murine lymph node tissue are well-established as a platform to study T cell motility.^41,54,55^ Otherwise, this system has seen limited use, particularly to study the response to polyclonal or antigen-specific stimulation.

At this time, unanswered questions regarding the viability, level of immune activation, and retention of function appear as potential obstacles to the broad adoption of live lymph node slices. To address this issue, here we describe a systematic evaluation of the procedures surrounding the slicing, handling, and analyzing of live murine lymph nodes in short-term cultures, towards establishing lymph node slices as a robust experimental platform. We comprehensively assess 24-hr viability, the extent of inflammation due to slicing, and retention of acute function. Finally, we validate the use of acute murine lymph node slices to quantify antigen-specific T cell responses ex vivo.

## Results

### Lymph node slices preserve spatial organization

We developed a protocol for slicing lymph node tissue that was informed by well-established procedures for slicing brain, another delicate tissue, and by prior work with lymphoid tissues.^41,45,54^ In brief, LNs were gently isolated from the animal, embedded in agarose for physical support, and sliced on a vibratome. LN slices were immediately immersed in culture media to rest until further processing or experimentation. A detailed experimental protocol is provided in the Methods. In this first section, we highlight some of the key aspects of working with lymph node slices prior to describing the optimization and validation of the method.

One of the primary reasons to work with intact tissue rather than cell culture is the preservation of spatial organization. Indeed, the structure of the lymph node was retained in these live, thick slices in the absence of fixation. Live tissue slices from naïve mice contained distinct B cell regions and lymphatic vasculature/vessels with a distribution that was consistent with in vivo and immuno-histochemical studies (Figure 1a).^25,39,45,56,57^ These geographical landmarks were readily visualized using widefield microscopy after live immunofluorescence staining.^43^ Ex vivo slices could also be used to visualize the distribution of draining antigen after in vivo vaccination, e.g. with rhodamine-conjugated ovalbumin (OVA) protein (Figure 1b-d). Both localization in individual cells (Figure 1b) and draining of soluble antigen via the lymphatic and sinus structure (Figure 1c-d) were visible without fixing the tissue.

**Figure 1:**
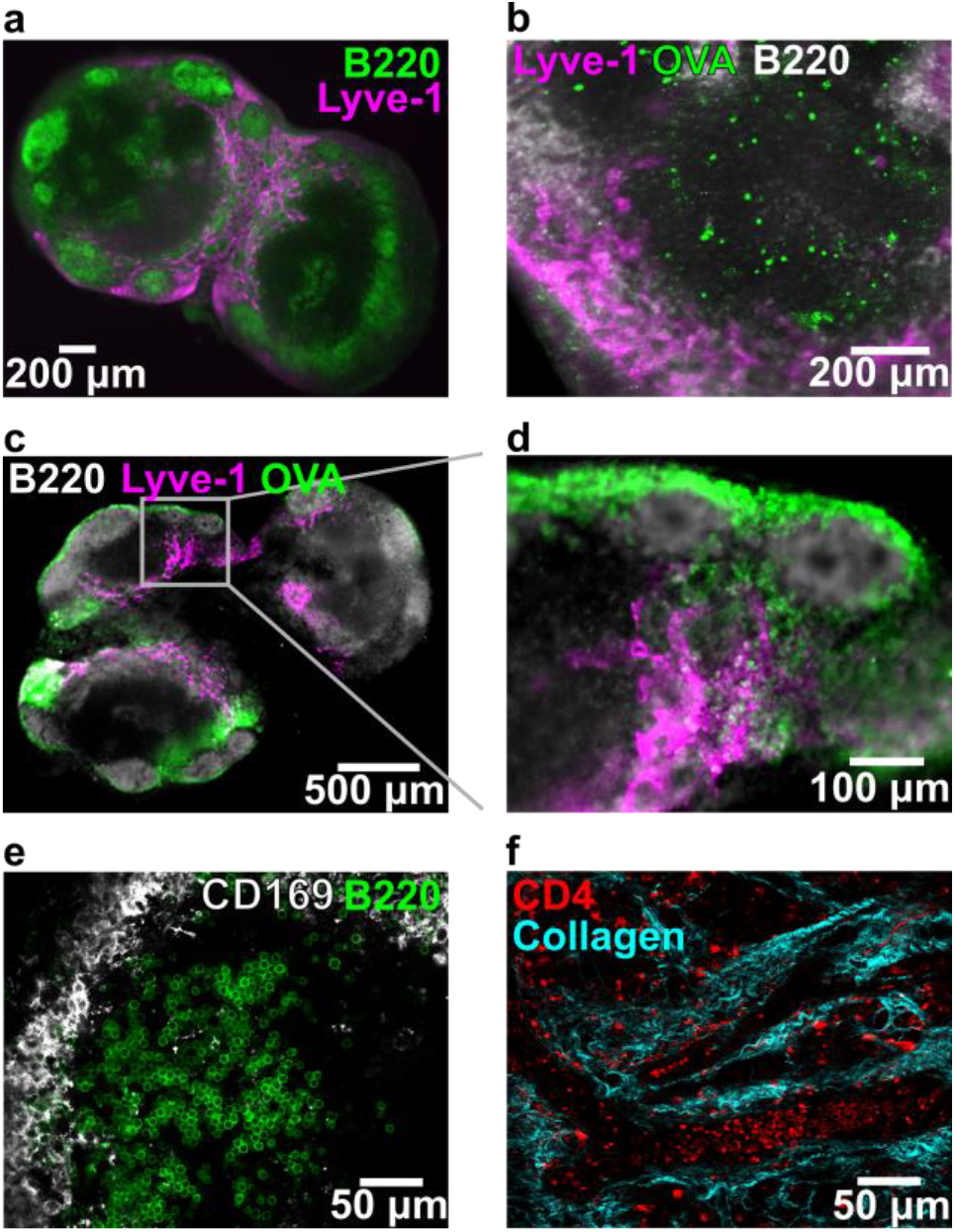
Key structural features remain intact in thick lymph node slices (iLN, aLN, bLN). (a) Slice labeled with anti-B220 (FITC, green) and anti-Lyve-1 (eFluor660, purple) revealed key structural features of the lymph node. Slice shown from a female C57Bl-6J mouse. (b-d) Slices from an OVA (rhodamine labelled, green)-immunized C57Bl/6-J mice, labeled with anti-B220 (grey) and anti-Lyve-1 (purple). (b) Rhodamine-OVA was visible inside of cells within the T cell rich (B220-dim) region of the lymph node 3 days after immunization. (c) Rhodamine-OVA was visible in the sinuses and lymphatics 1-day after immunization; panel (d) shows inset. (e) High-definition image collected by confocal microscopy. Slice labelled with anti-B220 (AlexaFluor 647, green) and anti-CD169 (AlexaFluor 594, grey). (f) Image of lymph node slice collected by two-photon microscopy, showing CD4 positive T cells (FITC-CD4 Fab’) within the collagen matrix (second harmonic imaging). Detailed methods for each panel are provided in the SI.

While the images described above were collected at low magnification, live tissue slices are also compatible with high resolution microscopy techniques. By using confocal microscopy, we were able to visualize individual cells with distinct morphologies within the slice, e.g. rounded B cells in a follicle and the network of macrophages surrounding its outer edge (Figure 1e). Second harmonic imaging of the collagen network within the LN slice (Figure 1f) highlighted the dense collagen network that persists throughout the lymph node, consistent with other examples of live two-photon imaging of the lymph node.^58–60^ These images highlight the potential for lymph node slices to reveal tissue organization.

### Lymph node slices reflect heterogeneity in organ composition

To enable quantitative cellular phenotyping and analysis of viability, we first determined that flow cytometry can be run on a single murine lymph node slice, similar to reports on single thymus slices.^45^ Manual counting indicated that each slice yielded, on average, (0.56 ± 0.16) ×10^6^ cells (n = 10 slices, mean ± std dev). The variability in cell number reflects the fact that the surface area varies between slices. For flow cytometry, single slices were crushed through a 70-μm filter to generate a cell suspension, in the same manner as is traditionally done for intact lymph nodes (Figure *2*a). We found that a single murine lymph node slice provided sufficient cell counts to collect flow cytometric data (Figure *2*b), and all subsequent analyses were performed on single slices unless otherwise noted. Live cells were identified using PI exclusion, and the remaining cells were separated into dead (PI^high^, DilC1^low^) and double positive (DP, PI^high^, DilC1^high^) populations according to signal from the mitochondrial membrane potential dye DilC1 (Figure *2* b). Live cells were phenotyped by surface markers for cell type (Figure *2*d-f).^61^

**Figure 2:**
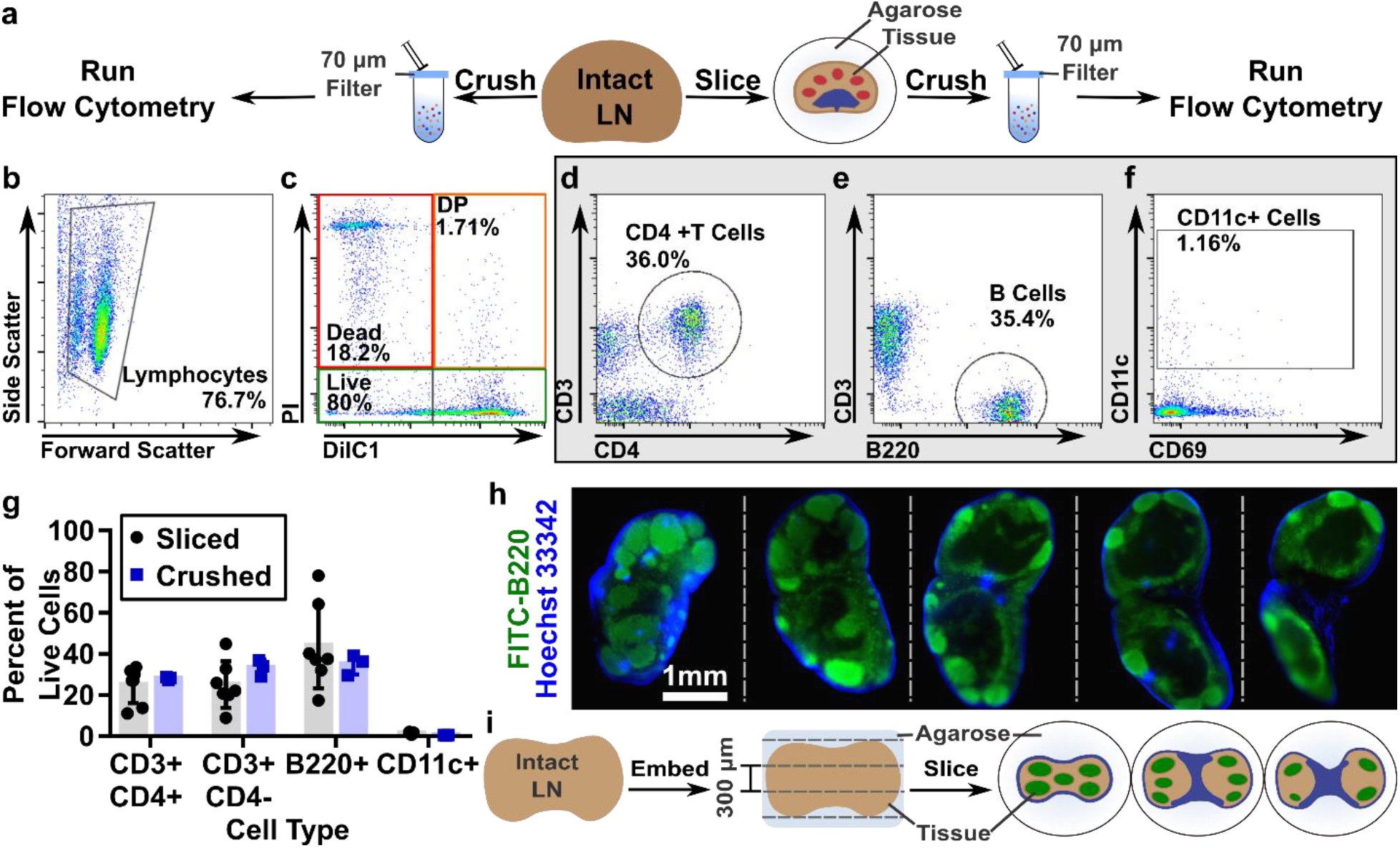
Lymph node slices are highly heterogenous. (a) Schematic showing experimental work flow: Intact LNs were either passed through a 70-µm filter or embedded in agarose, sliced, cultured, and then passed through a filter for flow cytometric analysis. (b-f) Representative flow plots from a single naïve slice showing scatter (b), viability (c), T cell, B cell, and CD11c+ cell phenotyping (d,e,f, respectively). DP indicates double positive (DilC1+PI+). (g) Average phenotypic distribution within individual lymph node slices compared to whole crushed lymph nodes. The average slice was equally distributed between B cells (44±21%) and CD3+ T cells (50±17%), with very few CD11c+ cells collected (1±0.4%); exact composition varied between slices. These data were not significantly different from whole crushed lymph nodes. Bars show mean ± standard deviation from N=7 sliced and N=3 crushed nodes pooled from iLNs, aLNs, and bLNs. (h) Serial 100-μm thick slices of a fixed lymph node labelled with FITC anti-B220 (green) and Hoechst 33342 (blue) detailing the heterogeneous cell distribution in the lymph node and how it changes with depth in the tissue. (i) Schematic representation of slicing the complex three-dimensional lymph node into 300-µm increments, which yields slices that are heterogeneous in terms of cell population and spatial distribution. B cell follicles shown in green; sinuses in blue.

On average, the cell suspension obtained from single C57Bl/6 lymph node slices matched that obtained from whole lymph nodes: 50% CD3+ T cells (51% CD4+, 49% CD4-), 44% B cells, and 1% CD11c+ cells (Figure *2*g). The low percentage of CD11c+ cells observed in both crushed and sliced samples is likely due to their adhesion to the extracellular matrix and failure to pass through the filter in preparation for flow cytometry.^39^ As expected, there was large heterogeneity in cellular composition between individual slices, as the slices reflect the complex 3-dimensional structure of this organ and the non-uniform distribution of cell types within it. In fact, immunofluorescence staining of thin (100-μm) serial slices of fixed lymph node tissue revealed significant heterogeneity from slice to slice in terms of both gross structural changes and cellular composition (Figure *2*h). The thicker, 300-µm slices of live tissue, are similarly heterogeneous, with their structure varying by depth in the organ (Figure *2*i). Thus, tissue slices of spatially organized organs may provide a means to quantify and assess variation in population function across the tissue, whereas methods that begin with tissue homogenization lose this information. Over dozens of experiments, we observed that variations in large scale tissue architecture between slices from the same organ exceeded the variations between the inguinal, axial and brachial lymph nodes. As the large-scale cellular architecture was similar for these three types of skin-draining lymph nodes, they were mixed for subsequent studies.

### Single 300-µm LN slices had similar overnight viability as lymphocyte suspensions

Next, we sought to optimize the conditions for lymph node slicing. First, we varied slice thickness to maximize the number of slices that can be collected from a single node while maintaining high viability. For reference, we compared cells collected from tissue slices to cells collected directly from intact lymph nodes by the conventional method of crushing through a filter (Figure *2*a). First, we determined the appropriate thickness for murine lymph node slices (Figure 3a); the minimal slice thickness for a given tissue depends on its mechanical strength, while an upper bound is set by its rate of oxygen consumption.^62^ Lymph node slices collected at 100 μm were usually torn, so this thickness was not considered further. 200-μm-thick slices were intact but sometimes mechanically distended (stretched); consistent with this, these slices were diminished in initial viability compared to 400-μm slices. There was no significant difference in initial or 24-hr viability between 300-μm and 400-μm slices, so 300 μm was selected to provide more slices per node. The percentage of live cells in slices was similar to that of cell culture suspensions over this time period (Figure 3a), indicating that the act of slicing did not significantly decrease the viability of the samples compared to crushing.

**Figure 3:**
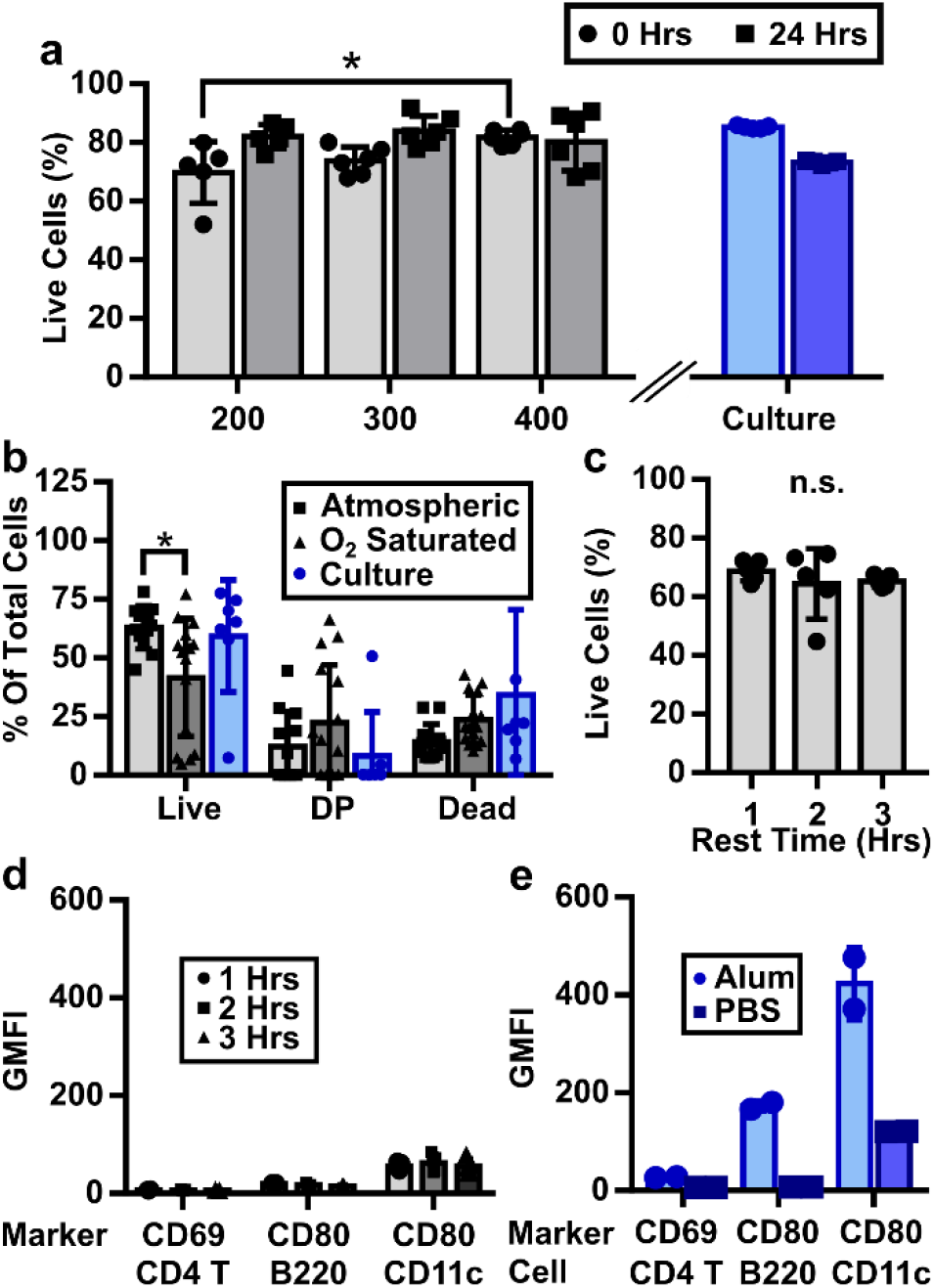
Optimizing parameters for LN slicing. (a) Tissue thickness had a slight effect on initial viability, as 200-μm thick slices were less viable than 400-μm thick slices. No significant differences were seen between thicknesses after 24 hours of culture. The viability of tissue slices was comparable to that of cell culture. (b) Slicing LNs in O_2_-saturated saline did not improve the viability of slices, as indicated by a decrease in the live and increased spread in the PI/DilC1 double positive (DP) populations. (c) Viability was unchanged over short recovery times. (d) The intensities of inflammation markers remained low over short recovery times and (e) were similar to cultures from crushed whole lymph nodes and much lower than those from lymphocyte cultures treated with alum in vitro as a positive control. Data collected from pooled iLNs, aLNs, and bLNs. 2-way ANOVA with multiple comparisons *p<0.05, n.s. p>0.05.

### Selection of slicing conditions to minimize activation markers

We aimed to select slicing conditions that minimized unintentional activation or alteration of the state of the lymph node, particularly the induction of rapid, non-specific inflammation due to mechanical damage from slicing. To do so, we varied the slicing conditions and analyzed viability and apoptotic markers, as well as the intensity of CD69 on CD4+ T cells and CD80 on B cells and CD11c-expressing cells, which include dendritic cells (DCs).^63–65^ We first considered the protein content and oxygenation of the media used during slicing. Inclusion of proteins in the chilled slicing media (i.e., addition of 2% v/v fetal bovine serum (FBS) to PBS) did not improve viability (Figure S1), so PBS was used for simplicity. Oxygenation of slicing media is essential for brain slices, and this convention has been propagated through many other tissue slicing protocols.^41,45,54,66,67^ However, lymph nodes, and many other tissues, are thought to be mildly hypoxic in vivo.^68^ We hypothesized that hyperoxia may not be needed during slicing of LN tissue. To test this hypothesis, we sliced tissue in PBS that was either bubbled or not with oxygen, after which the slices were rested in a cell culture incubator in complete media. Slices collected in oxygen-saturated PBS showed a small but significant decrease in the live population compared to those sliced under atmospheric conditions and trended towards a greater DP population (Figure 3b; tissues analyzed 1 hr post slicing). CD80 expression was also increased on CD11c positive cells from these slices (Figure S2). From these data we concluded that an O_2_-saturated environment during slicing did not improve lymph node slice viability, and so for simplicity all slices were collected in 1x PBS without oxygen bubbling. We note that these results were collected on skin-draining lymph nodes, and we cannot exclude the possibility that lymph nodes from other areas of the body may require different handling.

Slices of many organs are “rested” for one or more hours after collection to allow any effects of cutting to dissipate,^30,35,69^ and we tested slices in this window for viability and upregulation of inflammatory markers. We found no significant difference in viability over a period of 1 – 3 hr after slicing (Figure 3c, Figure S3), nor any increase in the fluorescence intensities of the activation markers CD69 on CD4+ T cells and CD80 on B cells and CD11c+ cells (Figure 3d, Figure S3). The CD69 and CD80 intensities from slices were comparable to lymphocytes collected directly from crushed nodes and cultured in 1x PBS, and were much lower than those from in vitro-activated lymphocytes that served as a positive control (Figure 3e, Figure S4). Based on these data, we determined that a 1-hour rest is sufficient post-slicing; shorter times may also be acceptable but were not tested. We speculate that the lack of measurable inflammation in response to the mechanical damage of slicing may be due to the rapid dilution of “danger signals” from the cut faces of the slice into the large volume of slicing media. In summary, lymph node slices collected in normoxic saline and rested for one hour displayed high viability and minimal markers of nonspecific activation.

### Inflammatory gene expression was low and similar between sliced and crushed lymph nodes

To further investigate the possibility of inflammation due to slicing, the expression of 84 key inflammatory genes (Table S1) was analyzed by RT-PCR array for lymph node tissue slices versus conventional lymphocyte cultures. The slices and cell suspensions were cultured overnight prior to analysis, to allow time for any delayed response or slow-acting inflammatory signals from the process of slicing, then mechanically dissociated directly into lysing buffer. Differential expression was determined by calculating the mean relative expression (sliced/crushed cultures) and setting a conservative threshold at one or two standard deviations from the mean. Consistent with the fact that these were samples from naïve animals, the majority of genes in the inflammatory gene array either were not expressed in either sample (27 genes) or were not differentially expressed between slice culture and cell culture (39 genes; Figure 4a), even with the least stringent threshold (1 std dev). Of the differentially expressed genes, 9 out of 18 were related to chemokines and their receptors (Figure 4b). We speculate that the increase in gene expression along the chemokine axis may be related to the preservation of the stromal cells and other matrix bound cells in the tissue slices, as these would have been removed by the filter when collecting lymphocyte suspensions.^38,70^ This observation remains to be explored. Based on the overall low levels of inflammatory gene expression in both tissue slices and cell suspensions, together with the low levels of activation markers observed by flow cytometry (Figure 3), we conclude that the process of slicing does not cause appreciable inflammation of the tissue. These data are consistent with results from tumor slices that found few changes in gene expression caused by the act of slicing.^71^

**Figure 4:**
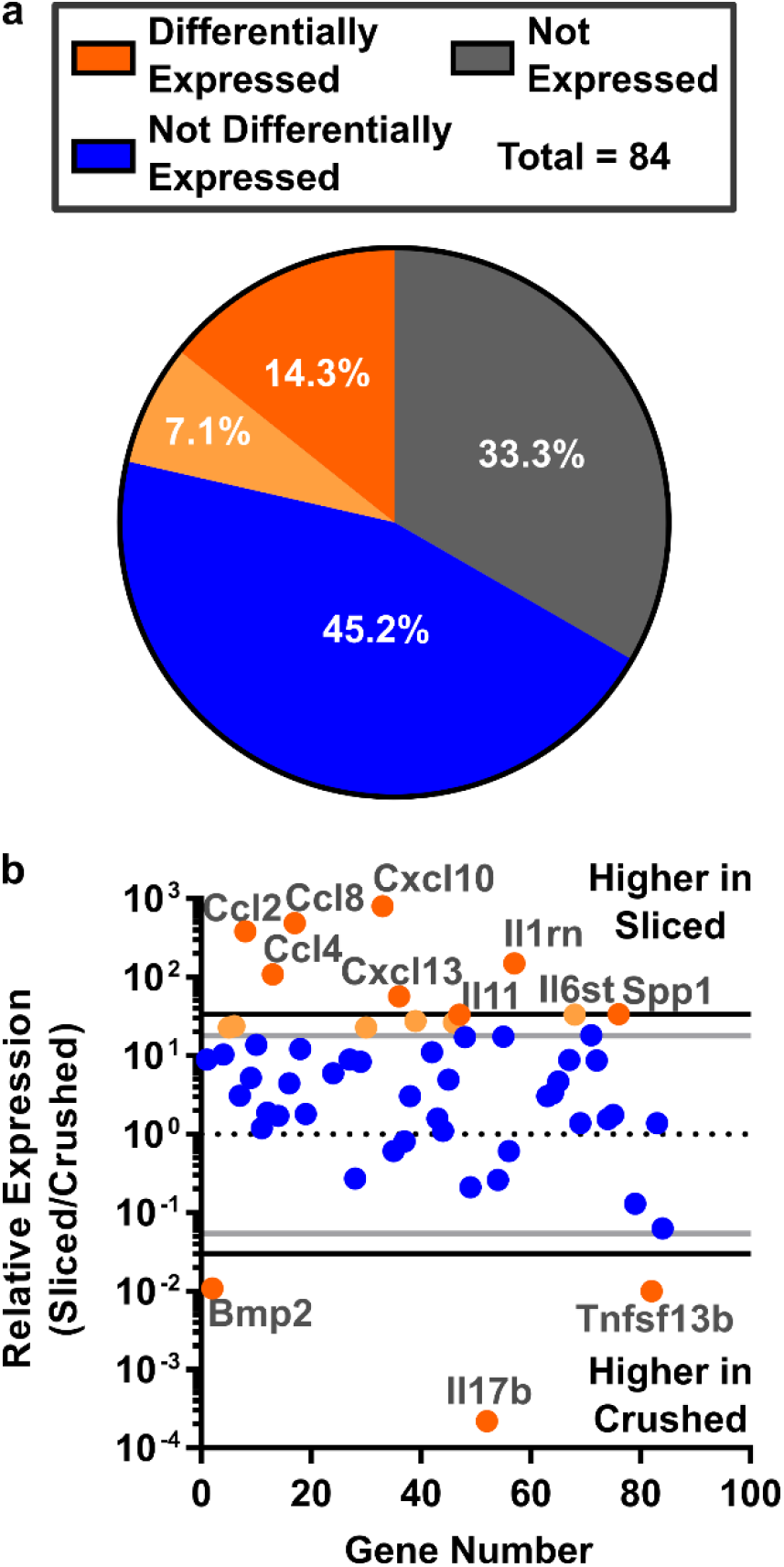
Comparable expression of inflammatory genes in slices and cell suspensions from naïve murine lymph nodes. (a) Of the 84 genes investigated, 33.3% were not expressed in either condition, 45.2% were expressed but not differentially expressed, and 14.3 (dark orange) to 21.4% (dark + light orange) were differentially expressed. Differential expression was determined by setting a threshold at 1 or 2 standard deviations from the mean relative expression of expressed genes (light and dark orange, respectively). (b) Expressed genes were categorized by the two cut-offs: 2 stdev (black lines, dark orange points) or 1 stdev (grey lines, light orange points). Seven genes were different between the thresholds. A full gene list is provided in Table S1. Gene expression was measured from pooled samples comprising 9 crushed or 9 sliced nodes (iLNs, aLNs, bLNs, 30 slices total) from N=3 mice.

### Lymph node slices processed whole-protein antigen and responded to cellular stimulation

An exciting application of lymph node slice culture is to measure the response of the intact tissue to ex vivo stimulation, with all cell types and structures present and correctly localized. We were particularly interested in the function of antigen-presenting cells, because appropriate antigen recognition is required to initiate adaptive immune responses. We tested the ability of antigen-presenting cells to process whole-protein antigen by incubating live slices with DQ-OVA, a modified form of ovalbumin that becomes fluorescent upon proteolytic cleavage. Repeated fluorescent imaging revealed time-dependent uptake and processing of the whole-protein antigen by cells in lymph node slices. Mean DQ-OVA intensity was significantly greater in live slices than in fixed slices after just two hours (Figure 5a). DQ-OVA signal followed a spatial distribution that was consistent with the sinuses and lymphatics (Figure 5b,c), reminiscent of the pattern observed for in vivo antigen drainage (Figure 1c-d), despite the bath method of delivery. Closer observation by 5-color confocal microscopy showed that the processed protein was visible inside F4/80+ macrophages, CD169+ macrophages, and Lyve-1+ lymphatic endothelial cells, but was mostly excluded from B220+ B cells (Figure 5d).^72,73^ Qualitatively, the largest fraction of processed protein at this time point appeared to be from CD169+ macrophages (Figure 5d, arrowheads, and Figure S5). Quantitative phenotyping of DQ-OVA positive cells would be best performed by disaggregating the tissue and performing flow cytometry; on the other hand, the multi-color imaging shown here offers complementary information by preserving the spatial organization of the cells and highlighting the preferential uptake near the sinuses and parafollicular zones.

**Figure 5:**
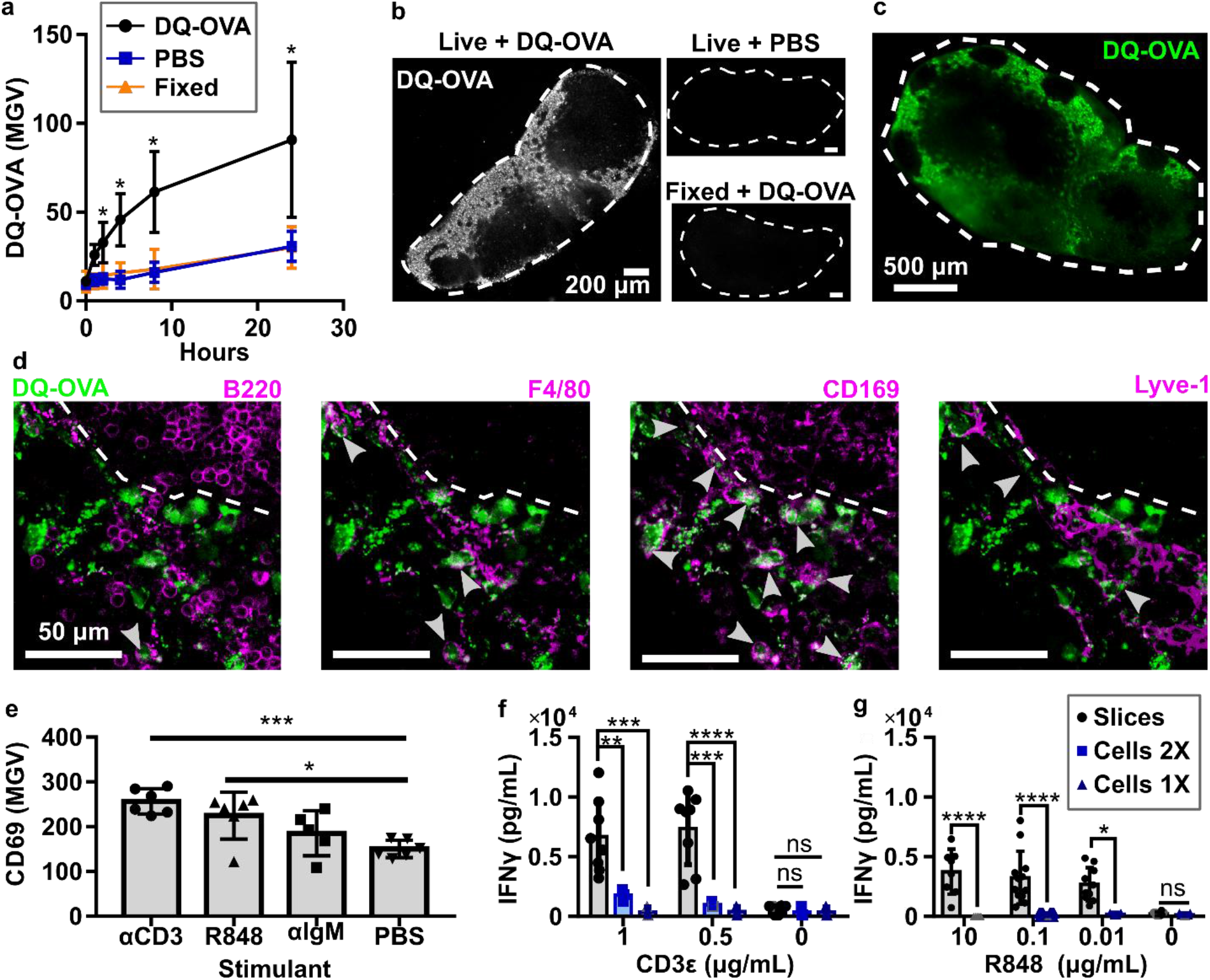
Slices processed protein antigen and responded to cellular stimulation. (a) Mean grey value (MGV) of DQ-OVA in lymph node slices, showing processing of this protein antigen only in live slices. Live slices incubated with 1x PBS and fixed slices incubated with DQ-OVA served as negative controls. 2-way ANOVA with multiple comparisons. N = 9 slices (iLNs, aLNs, bLNs), *p<0.05. (b) Representative images of slices from (a) after 4 hours of culture. Slices outlined with dashed white line from brightfield images (not shown). (c) Low-magnification, widefield image of DQ-OVA processed within a live slice after four hours. (d) Confocal images of a representative area of the slice shown in panel (c). Each panel includes DQ-OVA (green) overlaid with the indicated co-stain (purple). Dashed line indicates edge of B cell follicle. Arrowheads indicate cells that appear to have processed DQ-OVA. (e) Mean pixel intensity of CD69, averaged within each slice, was increased after 24-hour stimulation with indicated reagents. One-way ANOVA with multiple comparisons. Each dot represents one slice. ***p=0.0005, *p=0.0110. (f,g) IFNγ secretion from slices and mixed lymphocyte culture after 20-hour direct (CD3ε, e) or indirect (R848, f) T cell stimulation. Lymphocyte concentration was matched to LN slice, 1X: 1.7 ×10^6^ cells/mL, 2X: 3.4 ×10^6^ cells/mL. (f) IFNγ secretion after 20-hr stimulation with CD3ε. (g) IFNγ secretion after 20-hr stimulation with R848. Grey points were set to the LOD for the plate; each dot represents one slice/cell culture. N = 6-12 slices and 3-4 cell cultures pooled from iLNs, aLNs, and bLNs. Mean ± Standard Deviation. 2-way ANOVA with multiple comparisons. *p=0.0102, **p=0.0029, ***p=.00001, ****p<0.0001, n.s. denotes p>0.05.

We next tested the extent to which lymph node slices were able to respond to the activation of T cells (anti-CD3, TCR engagement), B cells (anti-IgM, BCR engagement) and APCs (R848, a TLR7 agonist) in overnight cultures.^74–76^ While anti-CD3 directly activates T cells by cross-linking the TCR, R848 acts on T cells indirectly by activating APCs to produce IL-12, which has a paracrine effect on nearby T cells.^77,78^ Rather than analyze cellular-level responses, which is best done by traditional cell culture and flow cytometry, we focused on readouts that reflect tissue-level responses and multi-cell interactions, such as upregulation of the lymphocyte activation marker CD69. Statistically significant increases in mean CD69 surface staining were induced by anti-CD3 and R848 but not anti-IgM (Figure *5*e). These stimuli generated qualitative differences in CD69 staining patterns within the tissue; anti-CD3 elicited diffuse CD69 signal in the T cell-rich paracortex, while other stimuli did not (Figure S6). Notably, anti-CD3 elicited these responses without inclusion of anti-CD28, suggesting that the co-stimulatory signal was adequately provided by the CD80/86 on APCs present within the tissue slice. Whether the T cell response is already maximized by anti-CD3, or could be enhanced by inclusion of anti-CD28, will be tested in future experiments.

We further tested the response of lymph node slices to ex vivo stimulation with anti-CD3 and R848 in terms of cytokine secretion, and directly compared the response of slices to lymphocyte suspensions at matched cell densities. As expected, both tissue slices and cell cultures responded to anti-CD3 with secretion of IFN-γ and IL-2, and unstimulated samples did not secrete measurable IFN-γ or IL-2 (Figure *5*f, Figure S7). Interestingly, we observed up to an 18-fold increase in IFN-γ but not IL-2 secretion in tissue slices compared to cell culture. Stimulation with R848 also resulted in significantly higher levels of IFN-γ production compared to lymphocyte culture (Figure *5*g). In fact, the lymphocyte response to these concentrations of R848 was very weak, possibly due to the relative scarcity of matrix-bound APCs in lymphocyte cell cultures.^39,79^ Splenocyte culture has a higher population of APCs compared to lymphocytes, and splenocytes did have a detectable response to R848, though still lower than lymph node slices (Figure S8).^80,81^ We speculate that the integrity of lymphocyte contacts with stromal cells and antigen-presenting cells (e.g. providing CD28 ligation), as well as the integrity of matrix-bound secreted factors, may play a role in differences between slices and cells. These data are consistent with prior reports that the cytokine profile from ex vivo-stimulated human tonsil and spleen slices differed from that of matched tonsil cell and splenocyte cultures.^50,53^ We conclude that lymph node slices can respond to ex vivo stimulation with both surface marker upregulation and cytokine secretion. Furthermore, differences between ex vivo and in vitro responses suggest that preserving the cell-cell and cell-ECM interactions that are seen in vivo may have a substantial impact on experimental results.

### Live lymph node slices responded to antigen-specific challenge

Finally, we tested the ability of lymph node slices to recall an antigen-specific response ex vivo. In vivo vaccination elicits a complex series of responses, including antigen trafficking and processing, cellular activation, and cytokine secretion, at specific times that reflect the ongoing development of the adaptive immune response. Using murine lymph node slices for antigen-specific responses is still rare,^82^ and to our knowledge has so far utilized peptide antigens and naïve tissue. Here we assessed the ability of ex vivo lymph node slices to report on the dynamic in vivo activities that occur in response to vaccination. Taking advantage of the OVA/OTII model antigen system, mice received CD4+ T cells expressing an OVA-specific transgenic TCR (OTII cells) intravenously, then were primed subcutaneously (s.c.) with either a model vaccine (alum and OVA protein) or a vehicle control (PBS) (Figure 6a). In this system we expected adjuvant-mediated and initial T cell responses between days 1 and 4, with a full T cell response by day 7.^83^ Alum-adjuvanted vaccines typically produce a Th2-skewed response, and we expected to see this polarization in our slice culture system in the form of IL-4 secretion.^84,85^

**Figure 6:**
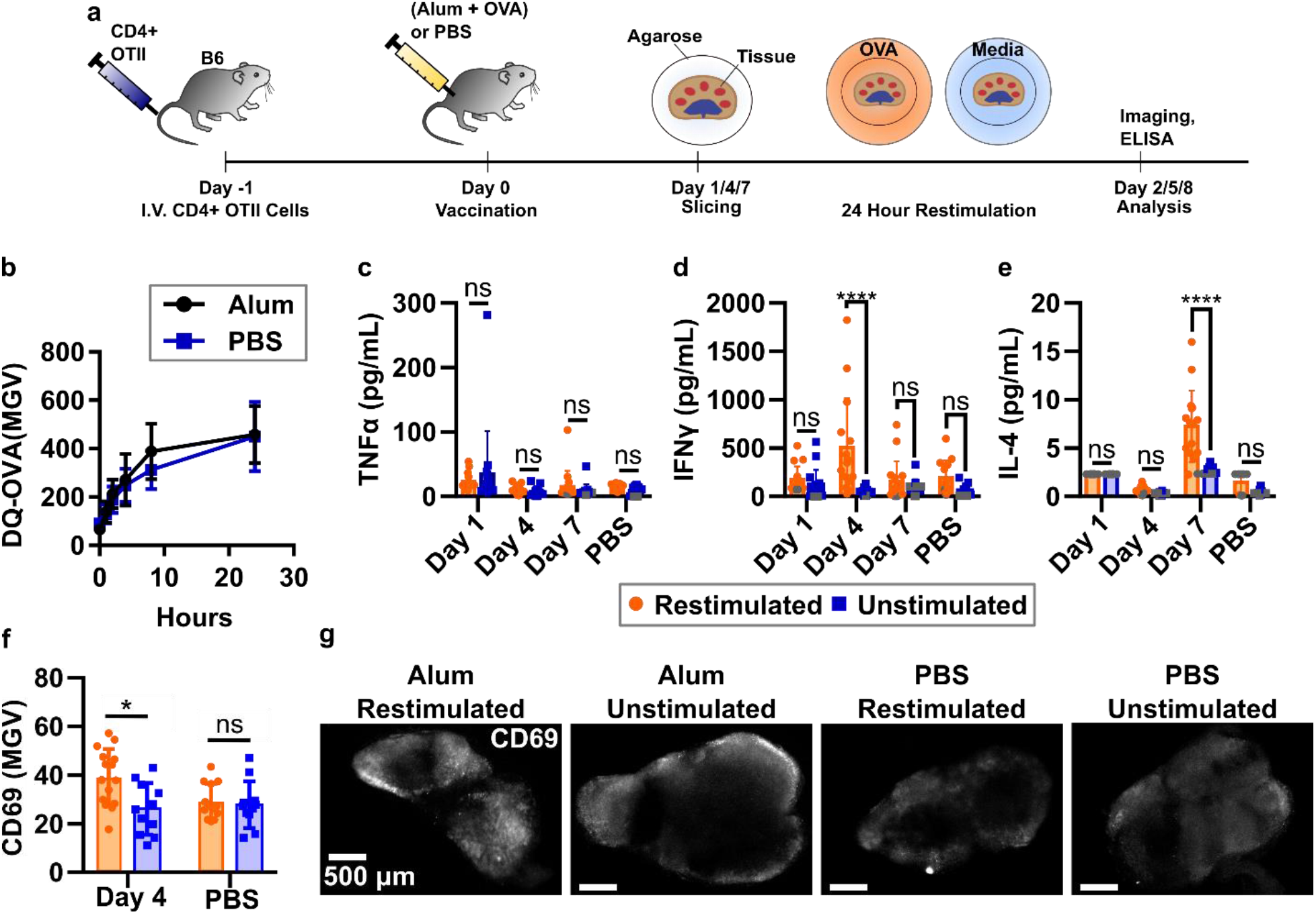
Lymph node slices showed antigen-specific stimulation after vaccination. (a) Schematic of experimental procedure. C57Bl/6 mice were given CD4+ OTII cells IV and rested one day before vaccination with either alum and OVA protein or PBS. Tissues were collected on 1, 4, and 7 days after vaccination and sliced. Slices were then cultured with or without OVA protein challenge for 24 hours. Cytokine analysis was completed by ELISA from surrounding media and slices were immunostained for activation markers. (b) Slices from mice vaccinated with alum+OVA four days prior processed OVA antigen at the same rate as those from unvaccinated mice. N=12 slices. (c-e) Quantification of cytokine secretion into the culture media after 24-hr culture with or without OVA. Antigen-specific IFN-γ response was detected on day 4; antigen-specific IL-2 response was detected on day 7. Dark grey points represent data that was set at the limit of detection for the plate. (f) Mean pixel intensity across an entire slice after live immunofluorescence of activation markers after vaccination with alum (day 4) or vehicle control (PBS). Slices were then exposed to either OVA protein or PBS as in cytokine studies. Each dot represents a single slice (iLNs, aLNs, bLNs) with bars indicating mean ± standard deviation. Two-way ANOVA with multiple comparisons. *p<0.05, **p<0.01, ****p<0.0001. (g) Representative images from experiment in (f). Slices were labeled with AF647-CD69. Scale bars all 500 μm.

Because activated APCs may decrease their phagocytic properties,^86^ we first tested whether cells in lymph node slices from vaccinated mice were able to process DQ-OVA on this time scale. Interestingly, DQ-OVA processing in these slices was unchanged compared to slices from control animals (Figure 6b). We speculate that the unaltered processing of DQ-OVA in the lymph node may be due to the presence of lymph-node resident DCs that were not activated by the s.c. alum injection. This result indicated that ex vivo incubation with antigen would still be effective in lymph node slices from vaccinated animals.

To quantify early and late responses to vaccination, lymph nodes were collected on days 1, 4, and 7 after vaccination, sliced, and cultured for 24-hr in the presence or absence of OVA protein in the media. TNFα secretion from the lymph node slice was not increased compared to control samples (naïve or unstimulated) at any time point (Figure 6c), consistent with other \reports for alum adjuvants.^84,87^ By day 4, a strong, antigen-specific IFN-γ response was detected in slices from vaccinated animals (Figure 6d). Slices from control animals did not secrete IFN-γ upon ex vivo culture with OVA, confirming that the response arose from vaccination. Consistent with the timing of the IFN-γ response, on day 4 we also observed a small but significant increase in CD69 immunofluorescence in vaccinated/restimulated slices compared to vaccinated/unstimulated slices and slices from unvaccinated animals (Figure 6f). CD69 upregulation was distributed throughout the slice (Figure 6g, Figure S9). No CD69 upregulation was detected on days 1 or 7 (not shown). As expected for an alum-adjuvanted vaccine,^83,85^ we also observed a statistically significant increase in antigen-specific IL-4 production in lymph node slices from vaccinated animals (Figure 6e). Interestingly, the peak days for IFN-γ and IL-4 differed (days 4 and 7, respectively), providing information about the kinetics of cytokine secretion during the in vivo immune response. Across all these data there was a wide spread of cytokine secretion between slices, which is most likely due to differing cellular composition. These slices were not pre-selected, as we were interested in measuring the robustness of this platform. However, slices in future experiments could be pre-screened prior to experimentation based on quantitative criteria such as the size of the T cell zone (e.g. by live immunofluorescence)^43^, to reduce variability in the results. In summary, these data provide compelling evidence that murine lymph node slices can mount antigen-specific responses ex vivo, including recall responses to intact protein antigens. In addition, these data show the potential of live lymph node tissue slices to provide a unique battery of multi-modal data, including antigen processing, cytokine secretion, and immunofluorescence labelling, to monitor an immune response as it occurs ex vivo.

## Discussion

The data in this paper lay out a set of best practices for slicing murine lymph nodes and maintaining them in 24-hr culture, and demonstrate that slices have the potential to be used to study T cell activation and antigen-specific responses ex vivo. Slices retained viability for 24 hours in culture and did not show signs of inflammation. Slices retained the spatial organization seen in vivo while making the tissue readily accessible for imaging and immunostaining, offering the potential to easily image cellular interactions and changes in surface marker expression during culture. Lymph node slices offered a cytokine response that differed in some cases from lymphocyte culture, consistent with prior reports for other lymphoid tissues.^50,53^ This difference may reflect the intact extracellular environment and cell-cell interactions that would be found in vivo. Most interestingly, this work provided evidence that lymph node slices could be used to report antigen-specific responses to vaccination, while offering the ability for multi-modal readout that combines imaging and traditional analysis such as ELISA and flow cytometry. Overall, this work lays the foundation for lymph node slices to serve as a controlled, ex vivo experimental platform in which to study the spatial organization and dynamics of the lymph node.

Traditionally, antigen presentation studies have been conducted by in vitro cell culture or by in vivo imaging, frequently using DCs pulsed with protein or peptide antigen.^56,59^ By using intact tissue ex vivo instead of co-culturing T cells with DCs in vitro, the contributions other cell types are retained, e.g. fibroblastic reticular cells (FRCs), with which T cells interact with at a higher frequency than with DCs.^88^ Furthermore, antigen-presenting cells were presumably present in same numbers and locations as they were in vivo. As ex vivo slices are well suited to image lymphocyte motility and cell-cell interactions that would otherwise be buried deep within the organ,^41,55,89,90^ they may be well suited to study T cell-DC interactions. Indeed, it is possible that dynamic events associated with antigen recognition could be achieved by overlaying pulsed DC cultures to the ex vivo slice platform, although overlays likely would not produce the fine segregation of DC phenotypes seen in vivo.^39,56,59^

The finding that cells in LN slices from vaccinated animals were able to perform the entire sequence from uptake and processing of protein antigens and to activation of cognate T cells extends the use of lymph node slices significantly. Here, the lymph node was primed in vivo, then challenged ex vivo with a protein antigen in a manner analogous to traditional vaccine challenge of lymphocyte suspensions. To our knowledge, the few prior reports of T cell activation in lymph node slices utilized either polyclonal stimuli (phytohaemagglutinin, etc.) or a peptide antigen for rapid activation.^43,52,82^ Unlike the protein antigen used here, which must be internalized and processed, peptide antigens may load onto any available MHC, thus triggering an initial T cell response even in naïve T cells. Here, cytokine secretion and CD69 upregulation were elicited only in lymph node slices from vaccinated animals, not from saline-control animals, indicating that the ex vivo response required prior in vivo activation by the vaccine. Thus, this work lays the foundation for the use of lymph node slices to analyze intercellular interactions between functional APCs and cognate effector T cells, particularly in the context of vaccination and also potentially in other settings where antigen-specific immunity develops.

Like any model system, ex vivo platforms have inherent features and limitations that impact experimental design. Ex vivo lymph node slices are characterized by heterogeneity, isolation from the organ, and route of exposure to the culture media that differs from the in vivo system:

1. *Heterogeneity*. The complex internal architecture of the lymph node results in each slice being distinct in both cellular distribution and available treatment surface area, despite all nodes being sliced along the transverse plane. This heterogeneity can lead to highly variable data sets, which can be addressed by either increasing sample size or by pre-selecting slices based on objective criteria, such as B220 intensity or slice area. Whether the heterogeneity is a drawback or a benefit depends on the question that one is asking. It may be possible to take advantage of this feature to tease out differences in responses in varying parts of the organ, instead of averaging over the entire population of the lymph node. While in principle, whole-organ explants could be coupled with two-photon imaging to provide similar information on regional variability, in practice the center of a whole-node explant is both difficult to image and likely to become hypoxic if the organ is larger than a few hundred microns in diameter. Drug stimulation and analysis of cytokine secretion are also limited by diffusion when working with whole-organ explants. Here, we preferred to keep the heterogeneous distribution of responses from lymph node slices as a representation of biological complexity rather than average or pool the data.
2. *Isolation*. Once the tissue has been removed from the body, no additional cells can be recruited from other areas, such as memory B cells from the bone marrow or circulating T cells from the blood stream.^91,92^ However, cells can be overlaid onto the slice ex vivo at defined concentrations and times, offering a powerful method to test the impact of specific cell types at known time points.^41,45^ Removal from the body also eliminates blood, lymphatic, and interstitial fluid flow, so shear stress is altered compared to what the organ is exposed to in vivo; this limitation is also present in standard lymphocyte cultures. Shear flow is known to affect the secretion of some chemokines (e.g. CCL21), as well as some lymphocyte functions.^93^ If needed, flow can be introduced through the tissue by gravity or perfusion culture,^47^ with control over flow rates to mimic different disease states in vivo and offer greater control of the experimental system. Microfluidics can offer a particularly controlled approach for both the perfusion of tissue and potentially long-term culture,^47,94–96^ though at the cost of additional complexity compared to a traditional cell culture plate.
3. *Route of access*. Drugs, cells, and detection reagents added to the culture media over a lymph node slice can enter the entire cut surface of the tissue, rather than being restricted to enter through the lymphatic or blood vasculature. This means that ex vivo slice treatment should not be used to report in vivo biodistribution, although uptake may still be somewhat selective based on regional cell activity (e.g. DQ-OVA uptake in Figure *5*b, and regional T cell homing reported previously^41^). On the other hand, this feature makes it possible to deliver stimuli to controlled locations rather than being limited by natural biodistribution mechanisms.^42,44,97^ Immunostaining of the live tissue by adding fluorescently labelled antibodies to the cut surface is straightforward, including for repeated quantitation to track changes in surface markers over time, which is challenging in vivo.^43^ Furthermore, in principle, antibodies against fixation-sensitive antigens should function better in these unfixed tissues than in traditional fixed sections, though this was not tested here.

Looking ahead, it will be useful to increase the longevity of the cultures to several days or weeks, to monitor an immune response from onset to completion ex vivo. Long term culture poses several challenges, including nutrient supplementation and oxygenation of the tissue, as well as retention of motile lymphocytes within the slices. Indeed, we have noted substantial egress of lymphocytes after 48 hours of sterile culture, similar to that observed in tonsil sections.^40^ Overcoming these challenges provides the opportunity to investigate the effect of individual environmental parameters, including nutrient concentrations and fluid flow profiles, on the basic biology of the lymph node, including both lymphocytes and stromal cells. As they currently stand, lymph node slices are able to provide valuable insight into short-term immune functions.

We are optimistic that live lymph node slices will provide a novel platform that will add to the immunologist’s tool box as a supplement to traditional experimental models. Slices provide a new angle of investigation based on spatially organized and dynamic cell-cell and cell-matrix interactions, coupled with the responses to well defined ex vivo stimulation. This makes them ideal to study the effect of both established drugs and novel compounds on the dynamics of lymph node tissue, an important step in generating and maintaining an adaptive immune response, especially in the context of vaccines. The enduring success of brain, lung, and tumor slices to study cellular and tissue-level events, pharmacological responses, and even response to damage and infection, indicates the wide array of translational utility for lymph node slices. Indeed, human tonsil slices already provide insight into the germinal center response and the response to viral infection.^40,52,53,98^ Tonsils are similar to other lymph nodes, with a greater frequency of germinal centers and increased T cell activation due to frequent exposure to pathogens.^99^ It seems likely that lymph node slices from transgenic animals and animal models of diseases such as cancer, autoimmunity and infection, will prove equally fruitful. Basic lymph node biology and physiology can also be compared across different microenvironments, e.g. by comparing gut-vs skin-draining lymph nodes. The approach described here for skin-draining lymph nodes has already been translated to murine mesenteric lymph nodes,^100^ and are expected to be compatible with other nodes as well. Comparing ex vivo function of nodes from different locations may highlight conserved structures and how the variations in these structures inform local immunity. Lymph node slices may also prove useful for monitoring the effect of the immune system on other organs, e.g. by co-culturing lymph node tissue with cells and tissues from elsewhere in the body, in a far simpler manner than is possible in vivo, while maintaining a high level of biological complexity.^47^ In summary, the methods for collecting and working with acute murine lymph node slices presented here provide a basis for the use of this model system to study dynamics and cellular interactions; applications may span the range of vaccine development, infectious disease, cancer immunity, and autoimmunity.

## Materials and Methods

### Generating lymph node tissue slices

All animal work was approved by the Institutional Animal Care and Use Committee at the University of Virginia under protocol #4042, and was conducted in compliance with guidelines the Office of Laboratory Animal Welfare at the National Institutes of Health (United States). Male and female C57BL/6 mice ages 6-12 weeks (Jackson Laboratory, USA) were housed in a vivarium and given water and food ad libitum. On the day of the experiment, animals were anesthetized with isoflurane followed by cervical dislocation. The inguinal, axial, and brachial lymph nodes (iLNs, aLNs, and bLNs respectively) were removed quickly and cleaned of any fat. It was critical to harvest the organ without deforming or puncturing it. Lymph nodes were placed in ice-cold DPBS without calcium or magnesium (Lonza, Walkersville MD, USA, #17-512F) supplemented with 2 % heat-inactivated fetal bovine serum (FBS, Gibco, Fisher Scientific, 100 % US Origin, 1500-500 Lot 106B14). Lymph nodes were embedded in 6 % w/v low melting point agarose (Lonza, Walkersville MD, USA) in 1X PBS. Agarose was melted in a microwave and allowed to cool until the temperature was comfortable in the hand. Further optimization showed that maintaining the melted agarose at 50 °C in a water bath after microwaving provided a more reproducible approach. Liquid agarose was poured into a 35 mm Petri dish, and lymph nodes were embedded in the liquid agarose close to the bottom of the dish. All lymph nodes were oriented to allow for the largest cross-section when slicing, i.e. a cut along the transverse plane. The dish was then rested at room temperature for approximately 2 minutes and allowed to harden on ice for the next 3 minutes. Once hardened, a 10 mm tissue punch (World Precision Instruments) was used to extract a section of agarose containing the lymph node. The block was inverted so the node was at the top of the section and glued onto a small mounting stage with Duro^®^ Super Glue (cyanoacrylate) and immediately submerged in a buffer tray containing ice-cold 1X PBS unless otherwise noted. Up to 6 lymph nodes were mounted on a single stage and sliced simultaneously.

A Leica VT1000S vibratome (Bannockburn, IL, USA) set to a speed of 90 (0.17 mm/s) and frequency of 3 (30 Hz) was used to slice 300-µm thick sections. A fan-shaped paint brush was used to remove the slices. Slices were immediately placed in a 6-well plate containing 3 mL per well of “complete RPMI”: RPMI (Lonza, 16-167F) supplemented with 10 % FBS (VWR, Seradigm USDA approved, 89510-186) 1x L-glutamine (Gibco Life Technologies, 25030-081), 50 U/mL Pen/Strep (Gibco), 50 µM beta-mercaptoethanol (Gibco, 21985-023), 1 mM sodium pyruvate (Hyclone, GE USA), 1x non-essential amino acids (Hyclone, SH30598.01), and 20 mM HEPES (VWR, 97064-362). Collection plates were equilibrated in a sterile cell culture incubator (37 °C with 5 % CO_2_) before slicing. Slices were rested in a sterile cell culture incubator for at least one hour prior to use. When sterile slices were required (for greater than 24 hours of culture) the vibratome was moved into a biosafety cabinet and isolated from the vibrations of the cabinet with a rubber-footed platform (Stand-Still Isolation Platform, Labconco). Sterile slices were collected as above with minor modifications (blade frequency set to 10 Hz).

### Activation of cell suspensions and tissue samples

Primary lymphocyte cell cultures were prepared by passing 6 peripheral nodes (axial, brachial, and inguinal) through a single 70-µm nylon mesh filter (Thermo Fisher, USA) with the rubber tip of the plunger from either a 1-or 3-mL syringe. Cells were plated in a 96-well cell-culture treated plate (Costar, VWR, USA) at a density of 1×10^6^ cells/mL in a 300 µL final volume. To obtain inflamed lymphocytes as positive controls, aluminum hydroxide gel adjuvant (Alhydrogel^®^, 10 mg/mL alum, Invivogen) was added to the wells for a final concentration of 1 mg/mL alum. Cells were cultured for 3.5 hours in a cell culture incubator (37 °C, 5% CO_2_) and prepared for flow as described below.

To compare activation of slices versus cell suspensions, peripheral lymph nodes (axial, brachial, inguinal) were randomly assigned to be sliced or crushed for lymphocyte culture. For the sliced condition, nodes were sliced 300 µm thick and each slice was placed into 500 µL complete media. For lymphocyte culture condition, nodes were crushed through a filter as described above. Lymphocyte suspensions were cultured in 500 µL aliquots at cell densities matched to tissue slice samples, where 1X culture was 1.7 ×10^6^ cells/mL, and 2X culture was 3.4 ×10^6^ cells/mL. Slices and lymphocyte cell culture were incubated for 20 hours at 37 °C, 5% CO_2_, with anti-mouse/human CD3ε (Biolegend, clone: 145-2C11, Purified grade) at 1, 0.5, or 0 µg/mL, with R848 (Resiquimod, InvivoGen, San Diego, CA) at 10 1, 0.1, or 0 µg/mL, or F(ab’)2 goat anti-mouse IgM (μ chain specific, Jackson ImmunoResearch) at 10 μg/mL.

### Flow cytometry

To prepare samples for flow cytometry, tissue slices were separated from the surrounding agarose through careful mechanical manipulation with a paint brush; individual tissue slices or groups of slices were then crushed through a 70-µm nylon mesh filter (Thermo Fisher, USA) using the rubber tip of a 1 or 3 mL syringe plunger to generate cell suspensions. Unsliced lymph nodes were similarly crushed through 70-µm filters, according to standard methods, for comparison. Cell suspensions were stained with Pacific Blue-B220, Brilliant Violet 421-CD3, Alexa Fluor 488-CD80, PE-CD11c, PE-Cy7-CD69, APC-Cy7-CD4 (all from Biolegend, USA, details provided in Table S2) and DilC1 (Thermo Fisher, USA). After staining, 2 µM propidium iodide (PI, Sigma Aldrich, USA) was added. Stained samples were washed and resuspended in 500 µL of 1x PBS with 2% FBS (flow buffer). Antibody compensation controls were run with OneComp eBeads™ (eBiosciences, USA) according to manufacturer protocol. Viability compensation controls, including PI and DilC1, were run on primary lymphocyte populations. PI controls were run with mixed live and killed cells; cells were killed with 35% ethanol for 10 minutes at room temperature. Live cells were stained with DilC1 for 30 minutes at 4 °C, washed and mixed with unstained live cells in a 1:1 ratio to act as a single stain compensation control. Stained suspensions were analyzed on a CyAn APD LX cytometer (Beckman Coulter, USA) unless otherwise noted. Analysis was completed using FlowJo 7 or FCS Express as noted.

### ELISA

Culture supernatant was collected and analyzed by sandwich ELISA for the cytokines IFNγ, IL-2, IL-4, and TNFα. A high-binding plate (Corning Costar 96 well ½ area, #3690; Fisher Scientific) was coated with 1 µg/mL anti-IFNγ XMG1.2, 1 µg/mL anti-IL-2 JES6-1A12, ELISA MAX capture anti-IL-4 (previous antibodies from Biolegend) or capture TNFα (R&D systems, cat: DY410-05) in PBS overnight at 4°C, then washed. All washing steps were performed in triplicate with 0.05% Tween-20 in PBS. Wells were blocked for 2 hours with 1% BSA and 0.05% Tween-20 (Fisher Scientific) in PBS (block solution). Serial dilutions of recombinant murine IFNγ, IL-2 (Peprotech, Rocky Hill, NJ), IL-4 (ELISA MAX standard, Biolegend) and TNFα (R&D Systems) were prepared in a 1:1 v/v mixture of block solution and complete media, and supernatant samples were diluted 1:1 v/v with block solution. Samples were added to the plate in duplicate and incubated for 2 hours, then washed. Biotinylated anti-IFNγ R46A2 (0.5 μg/mL), anti-IL-2 JES6-5H4 (1 μg/mL), ELISA MAX detection anti-IL-4 (Biolegend), or detection TNFα (R&D Systems) were prepared in blocking solution and added to the plate. Avidin-HRP (1X) (Fisher Scientific) in blocking solution was added to the plate and incubated for 30 minutes, then washed. Plates were developed using TMB substrate (Fisher Scientific), stopped with 1M sulfuric acid (Fisher Scientific), and absorbance values were read at 450 nm on a plate reader (CLARIOstar; BMG LabTech, Cary, NC). To determine concentration of sample solutions, calibration curves were fit in GraphPad Prism 6 with a sigmoidal 4 parameter curve (Eq. (1)), where X is concentration, Y is absorbance, min and max are the plateaus of the sigmoidal curve on the Y axis, and HillSlope describes the steepness of the slope.

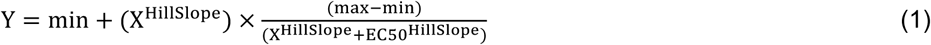

Limit of Detection (LOD) was calculated from the average of the blank + 3x standard deviation of the blank.

### Inflammatory Gene Expression Array

Axial, brachial, and inguinal lymph nodes from three mice were mixed and randomly distributed into two groups: 9 nodes for slicing and 9 nodes for cell suspensions. Approximately 30 slices were collected as described above and cultured individually at 37 °C with 5% CO_2_ overnight. Meanwhile, lymphocyte suspensions from whole nodes were generated by passing the lymph nodes through a 70-µm filter. The lymphocytes were pooled and resuspended at 0.86 ×10^6^ cells/mL (mean cellular density matched to the lymph node slices) then cultured overnight. After the overnight culture period all samples were flash frozen and stored at −80 °C until RNA could be isolated.

RNA was isolated using a RNeasy Mini Kit according to manufacturer instructions (Qiagen, USA). Briefly, pooled tissue samples (intact slices or cell culture suspensions) were homogenized in lysate buffer; cells were vortexed in lysate buffer and passed through a 20-gauge needle to generate a homogenized sample. Lysates were mixed with 70% ethanol and filtered according to manufacturer recommendations to obtain genetic material. To remove genomic DNA from the sample, 1 μg RNA was added to 1 U/µL DNase (Invitrogen, USA) in DNase reaction buffer. The digestion was run for 15 min at room temperature and stopped with 25 mM EDTA and heated to 65 °C for 10 min. An Accuris qMax cDNA synthesis kit was used to generate the cDNA. Reaction buffer, qMax reverse transcriptase, RNA and water were incubated at 42 °C for 30 minutes. The reaction was stopped by heating to 85 °C for 10 minutes.

A RT^2^ profiler array for mouse inflammatory cytokines and receptors (Qiagen, USA) was used according to manufacturer recommendations to measure the expression of 84 inflammatory genes (Table S1). SYBR Green was used as the reporter and the reaction was run for 40 cycles on a QuantStudio 6 PCR instrument (Thermo Fisher, USA). Genes that were detected on or after cycle 35 were considered not expressed. Of the expressed genes, the average relative expression was determined based on the average expression of 5 housekeeping genes (Table S1) Because all samples in each group were pooled for analysis, two cut-offs were defined for significant differential expression, two and three standard deviations from the mean differential expression.

### DQ-OVA culture of lymph node slices

Slices were collected as above and randomly assigned to live culture or fixation. Live slices were cultured with ovalbumin (OVA) protein solution, consisting of 1 μg/mL DQ-OVA (Thermo Fisher, USA) plus 9 μg/mL purified OVA (InvivoGen, USA) in 500 μL supplemented RPMI, or vehicle control in 500 μL supplemented RPMI. Killed control slices were fixed in formalin (4% formaldehyde, Protocol, USA) for 1 hour at 37 °C with 5% CO_2_, then incubated with OVA protein solution. Slices were incubated for 24 hours at 37 °C, 5% CO_2_, and images were collected at 1,2,4,8 and 24 hours.

### Antigen-specific interactions in lymph node slices

Spleens were collected from Rag2/OT-II female mice (Taconic Biosciences) aged 6-10 weeks following isoflurane anesthesia and cervical dislocation. Splenocytes were isolated using a 70-µm pore size nylon filter (Fisher Scientific, USA), and the filter was washed with sterile 1x phosphate buffer saline (PBS) supplemented with 2% v/v fetal bovine serum (FBS, VWR, USA). Cell density was determined through trypan blue exclusion. Using a CD4+ T cell enrichment kit (StemCell Technologies, USA), CD4+ T cells were isolated from bulk splenocytes by negative selection. OTII CD4+ T cells (0.5×10^6^ cells: 200 μL at 2.5×10^6^ cells/mL) were intravenously injected into 8 female C57Bl/6 mice. The following day, the C57Bl/6 mice were vaccinated with 50 µg of OVA protein in either 200 µL of Alum 50:50 v/v PBS or PBS alone. Vaccinated mice were humanely euthanized on days 1,4 and 7 after vaccination, and lymph nodes were harvested and sliced. Slices were cultured overnight in complete media supplemented with 10 µg/mL OVA protein (Invivogen) or PBS. After overnight culture, the supernatant was collected for cytokine analysis using ELISA and the slices immunostained and imaged.

### Immunofluorescent staining and imaging of lymph node slices

Slices were stained according to previously published procedures.^43^ Briefly, slices were placed on a Parafilm^®^ covered surface and a washer was placed on top. Samples were treated with blocking solution (anti-CD16/32) for 20 minutes in a cell culture incubator. Antibody cocktail was added to the blocking solution and samples were incubated for an additional 1 hour. Slices were then washed in sterile 1x PBS for at least 30 minutes in a cell culture incubator. Where noted, antibody Fab’ fragments were produced in house by pepsin fragmentation, as previously reported.^101^

Unless otherwise noted, imaging was performed on a Zeiss AxioZoom upright microscope with a PlanNeoFluor Z 1x/0.25 FWD 56mm objective, Axiocam 506 mono camera and HXP 200 C metal halide lamp (Zeiss Microscopy, Germany). Images were collected with Zeiss Filter Sets 38 HE (Ex: 470/40, Em: 525/50), 43 HE (Ex: 550/25, Em: 605/70); 64 HE (Ex: 587/25, Em: 647/70); and 50 (Ex: 640/30, Em: 690/50). Confocal microscopy was performed on a Nikon A1Rsi confocal upright microscope, using a 487 and 638 nm lasers with 525/50 and 685/70 nm GaAsP detectors respectively. Images were collected with a 40x/0.45NA Plan Apo NIR WD objective. Two-photon microscopy and second harmonic imaging were performed in the W.M. Keck Center for Cellular Imaging (University of Virginia) on an Axiovert200 MOT inverted microscope with an LSM510 scan head (Zeiss, Germany). Image was collected with 60x/1.20 WD objective. Image analysis was completed using ImageJ software 1.48v.^102^

## Supplementary Information

Supplemental methods for Figure 1, supplemental figures, and supplemental tables for gene expression data and antibodies used.

## Acknowledgements

The authors thank the Hartwell Foundation for generously supporting this work. Research reported in this publication was also supported by the National Institute of Allergy and Infectious Diseases under Award Number R01AI131723 through the National Institutes of Health. M. C. Belanger was supported in part by the Immunology Training Grant at the University of Virginia (NIH, 5T32AI007496-23). The content is solely the responsibility of the authors and does not necessarily represent the official views of the National Institutes of Health. Finally, the authors acknowledge the Flow Cytometry Core and the W.M. Keck center for cellular imaging at the University of Virginia for their technical expertise.

## Author Contributions

M.C.B. designed the research and was involved in the majority of data collection. A.G.B and M.A.C. designed and performed the antigen-specific response experiment in Figure 6. A.W.L.K designed and performed the slice vs cells activation experiments in Figure 5. P.A. and B.D.G. collected many of the immunofluorescent images used in Figure 1 and Figure 2. S.J.M. provided expertise for the inflammatory gene array experiment in Figure 4. A.E.R. designed and collected data for Figures 2 and 3 and drafted the introduction. J.R.L. and R.R.P. aided throughout in the design of the research and analysis and interpretation of data, and R.R.P edited the manuscript. M.C.B. took the lead in drafting the manuscript with contributions or revisions from all authors.

## FOR TABLE OF CONTENTS USE ONLY

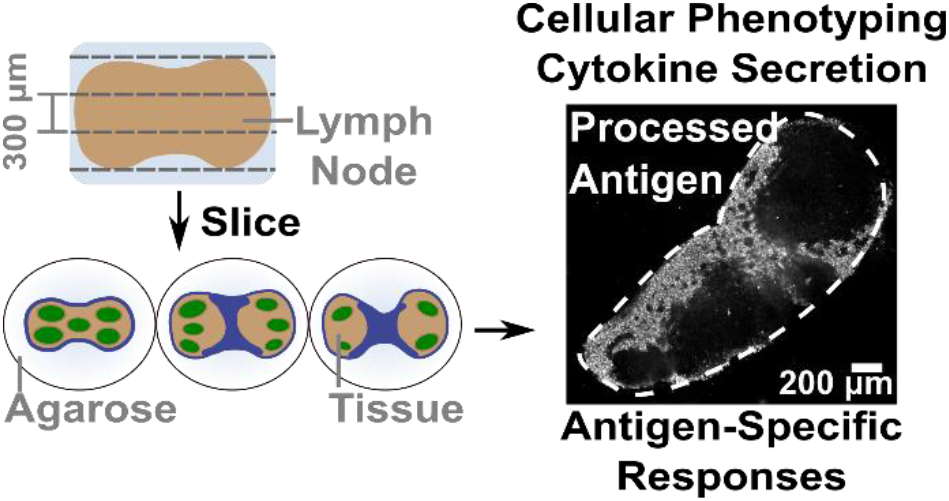

This article describes best practices and validation of the use of live, thick slices of murine lymph nodes in acute culture for ex vivo study of immune responses, including phenotyping, responses to molecular stimuli, antigen processing, and antigen-specific responses.

