## Supplemental Information for "Acute lymph node slices are a functional model system to study immunity ex vivo"

**Contents**


- Supplemental Methods
- Supplemental Figures S1 - S9
- Supplementa Tables S1 - S2
- Supporting References

**Supplemental Methods**

**Methods for experiments shown in Figure 1.**

**(a)** Slices were collected from a naïve C57Bl/6 female mouse aged 6-8 weeks (Jackson Laboratories, USA) and immunostained according to published procedures.[1] Slices were stained with FITC-B220 and eFluor 660-Lyve1. Image was collected on an AxioZoom (Zeiss, Germany).

**(b)** Naïve C57Bl/6 female mice aged 6-8 weeks were immunized subcutaneously with 50 µg/mL rhodamine-ovalbumin (OVA) protein. Mice were euthanized after 3 days and slices from draining lymph nodes were collected. Slices were immunostained according to published procedures with FITC-B220 and eFluor 660-Lyve1.[1] Image was collected on an AxioZoom (Zeiss, Germany).

**(c/d)** Naïve C57Bl/6 female mice aged 6-8 weeks were immunized subcutaneously with 50 µg/mL rhodamine-ovalbumin (OVA) protein. Mice were euthanized after 24 hours and slices from draining lymph nodes were collected. Slices were immunostained according to published procedures with FITC-B220 and eFluor 660-Lyve1.[1] Image was collected on an AxioZoom (Zeiss, Germany).

**(e)** Slices were collected from a naïve C57Bl/6 female mouse aged 6-8 weeks (Jackson Laboratories, USA) and immunostained according to published procedure with AlexaFluor-647-B220 and Alexa-Fluor 594-CD169.[1] Confocal microscopy was performed on a Nikon A1Rsi confocal upright microscope, using a 561 nm and 638 nm lasers with 600/50 and 685/70 nm GaAsP detectors respectively. Images were collected with a 40x/0.45NA Plan Apo NIR WD objective.

**(f)** Slices were collected from a naïve C57Bl/6 female mouse aged 6-8 weeks (Jackson Laboratories, USA) and stained with FITC-CD4 Fab' as previously reported.[1] The antibody fragment was generated by pepsin cleavage and reduction and conjugated to fluorescein succinimidyl ester in-house.[2] After staining, slices were fixed in formalin for 30 minutes and imaged. Two-photon microscopy and second harmonic imaging was performed in the W.M. Keck Center for Cellular Imaging (University of Virginia) on a Axiovert200 MOT inverted microscope with an LSM510 scan head (Zeiss, Germany). Image was collected with 60x/1.20 WD objective.

**Supplemental Figures and Tables**

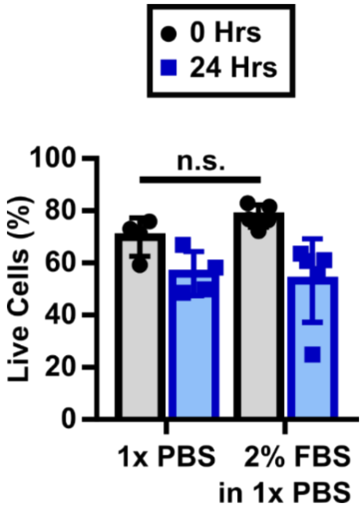


**Figure S1: Inclusion of serum in slicing media did not increase viability.** Slices were collected in either ice cold 1x PBS and PBS supplemented with 2% FBS. There was no significant differences between these conditions at either time point. Two-way ANOVA with multiple comparisons; ns denotes p>0.05.

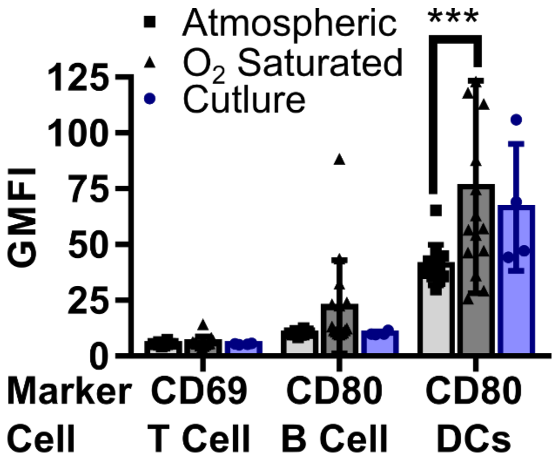


**Figure S2: Slicing tissue in an oxygen-saturated environment resulted in increased surface expression of inflammatory markers.** Activation markers on indicated cell types in slices collected under atmospheric or $O_2$-saturated conditions, compared to traditional lymphocyte culture. Slices collected under atmospheric conditions had statistically significant lower expression of CD80 on CD11c+ cells compared to slices collected in an oxygen-saturated environment but not significantly lower than lymphocyte culture. While trending higher the expression of CD80 on B cells in slices collected in an oxygen-saturated environment it was not significantly different compared to the other conditions. Each dot represents a single slice from skin-draining lymph nodes or cell culture well. ***p=0.0003, 2-way ANOVA with multiple comparisons.

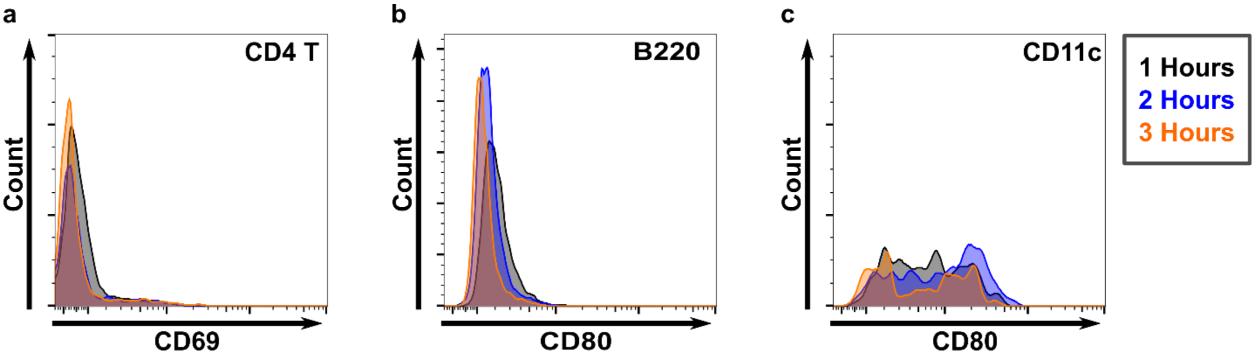


**Figure S3: Recovery times of greater than 1 hour did not significantly affect activation marker expression.** Slices of naïve lymph node tissue were collected and allowed to rest for 1-3 hours, and surface marker expression was determined by single-slice flow cytometry. Representative histograms (N=4-5 slices per condition) of (a) CD69 expression on CD4 T cells, (b) CD80 expression on B220 cells, and (c) CD80 expression on CD11c cells. Indicated populations gated as in Figure 2. There are no observable differences in histogram profiles across this time scale.

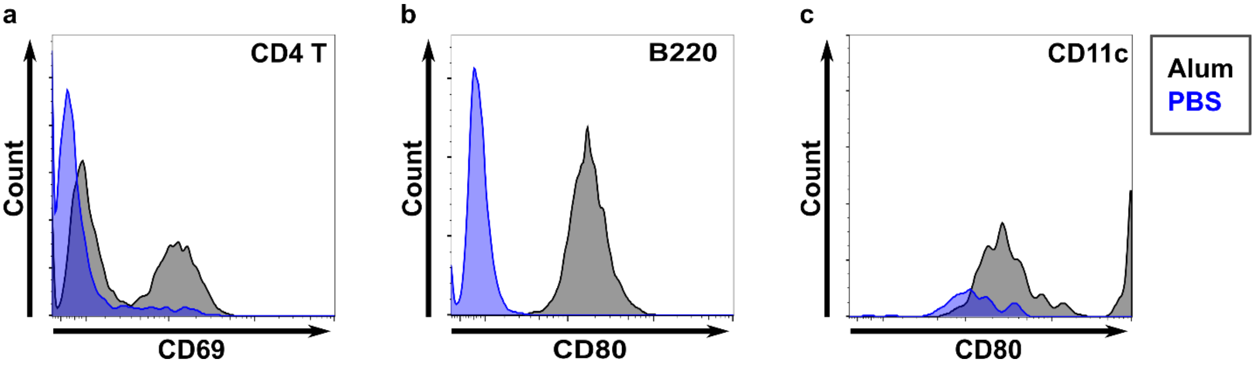


**Figure S4: Lymphocyte cultures responded to alum stimulation.** Lymphocyte cultures were stimulated with 1 mg/mL alum or 1x PBS for 3.5 hours in vitro and surface marker expression was quantified by flow cytometry. Representative histograms (N=2 samples) of (a) CD69 expression on CD4 T cells, (b) CD80 expression on B220 cells, and (c) CD80 expression on CD11c cells. Indicated populations gated as in Figure 2. In all cases, the alum-activated cultures expressed higher levels of activation markers on their surface. These cells acted as a positive control for activation marker expression.

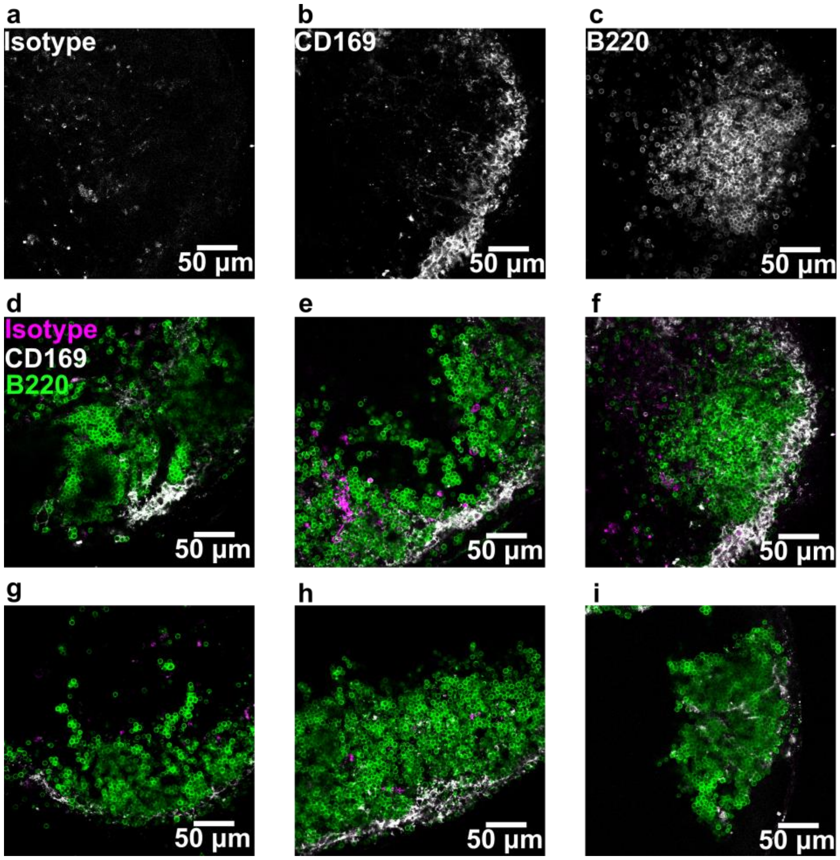


**Figure S5. CD169 and isotype control staining in B cell follicles in murine lymph node slices.** (a-c) Individual fluorescence channels from an isotype control immunostain (Rat IgG 2a, kappa), anti-CD69, and anti-B220. The individal channels clearly show the specifity of the CD169 signal in comparison to the isotype. (d-i) Fluorescence images of six different lymph node slices, with the B220 (green), CD169 (white) and isotype (purple) overlaid. Panel (f) shows the overlay of the data from panels (a-c). Much of the staining of CD169+ cells was found in the subcapsular sinus on the outter edge of the slices rather than throughout the follicle or alone its interior edge.

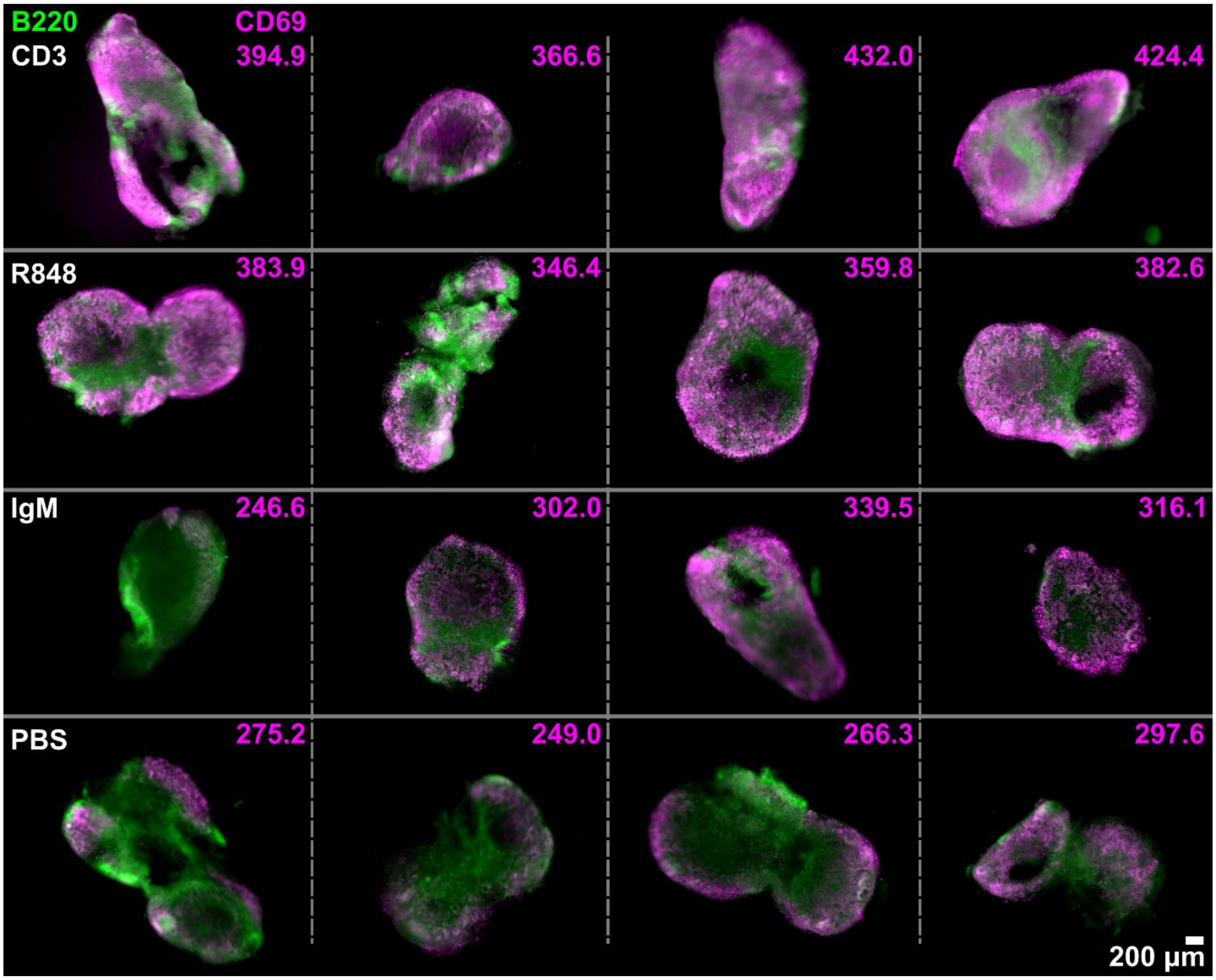


**Figure S6: Slices responded to stimulation by upregulating surface expression of inflammation markers**. B220 (green) and CD69 (purple) immunofluorescence in representative lymph node after ex vivo stimulation with anti-CD3 (row 1), R848 (row 2), anti-IgM (row 3), and PBS (row 4). Mean grey value for the CD69 channel is reported in the upper right corner of each image. The PBS control samples had visible CD69 staining around the edges of the tissue, due to either natural high expression in naïve lymph nodes or off target/Fc-mediated binding fo the antibody in these regions. Slices stimulated with anti-CD3 or R848 both had elevated, somewhat punctate expression within cortical regions of the lymph node, which are rich in both B cells and T cells, and CD3-stimulated slices also had diffuse CD69 signal in the T cell-rich center. Anti-IgM stimulation resulted in a uniform staining pattern but with much lower average intensity over the entire slice.

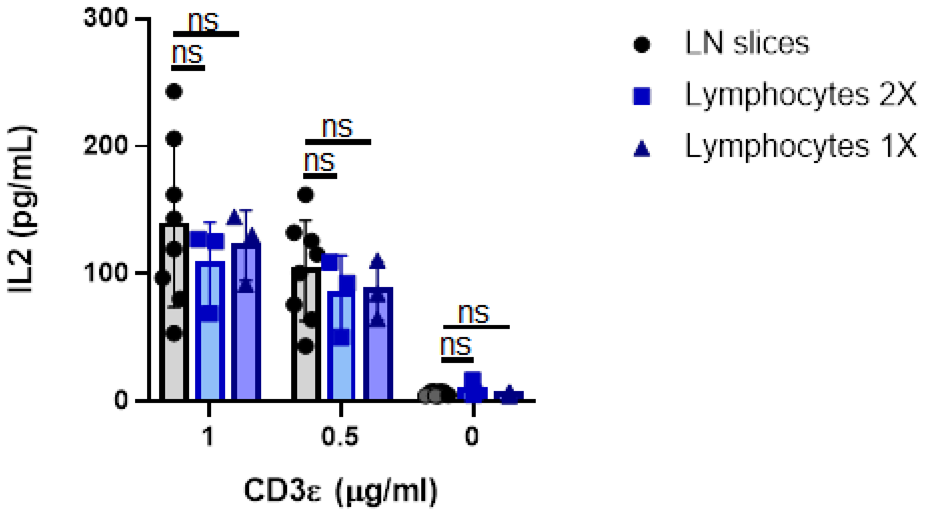


**Figure S7: Lymph node slices produced similar levels of IL-2 as mixed lymphocyte culture after anti-CD3 stimulation.** Lymphocyte concentration is matched to LN slice, 1X: 1.7 x$10^6$ cells/mL, 2X: 3.4 x$10^6$ cells/mL. Grey points indicate data that were set to the limit of detection. LOD = Avg of blank + 3xStd Dev of blank. Each dot represents one slice/cell culture. N = slices: 6-12; cell cultures: 3-4 per condition. Error bars denote standard deviation. 2-way ANOVA with Sidaks multiple comparisons; ns denotes p>0.05.

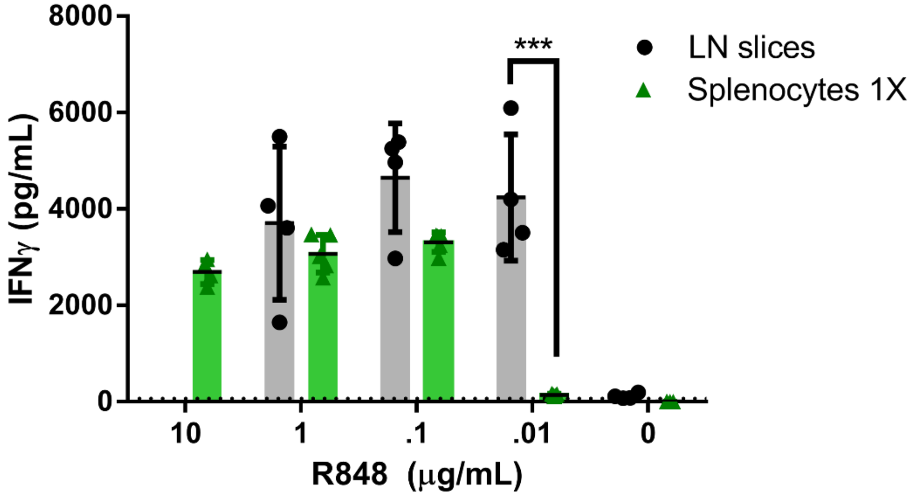


**Figure S8: Lymph node slices and mixed splenocyte culture responded to R848 stimulation.** R848-stimulated lymph node slices had greater IFNγsecretion than splenocytes at matching cell concentrations, most notably at low concentrations of R848 (0.1 µg/mL). N=4 slices per condition. Dotted line indicates LOD for the plate. Error bars denote standard deviation. 2-way ANOVA analysis with Sidaks multiple comparisons; ***p=.00001

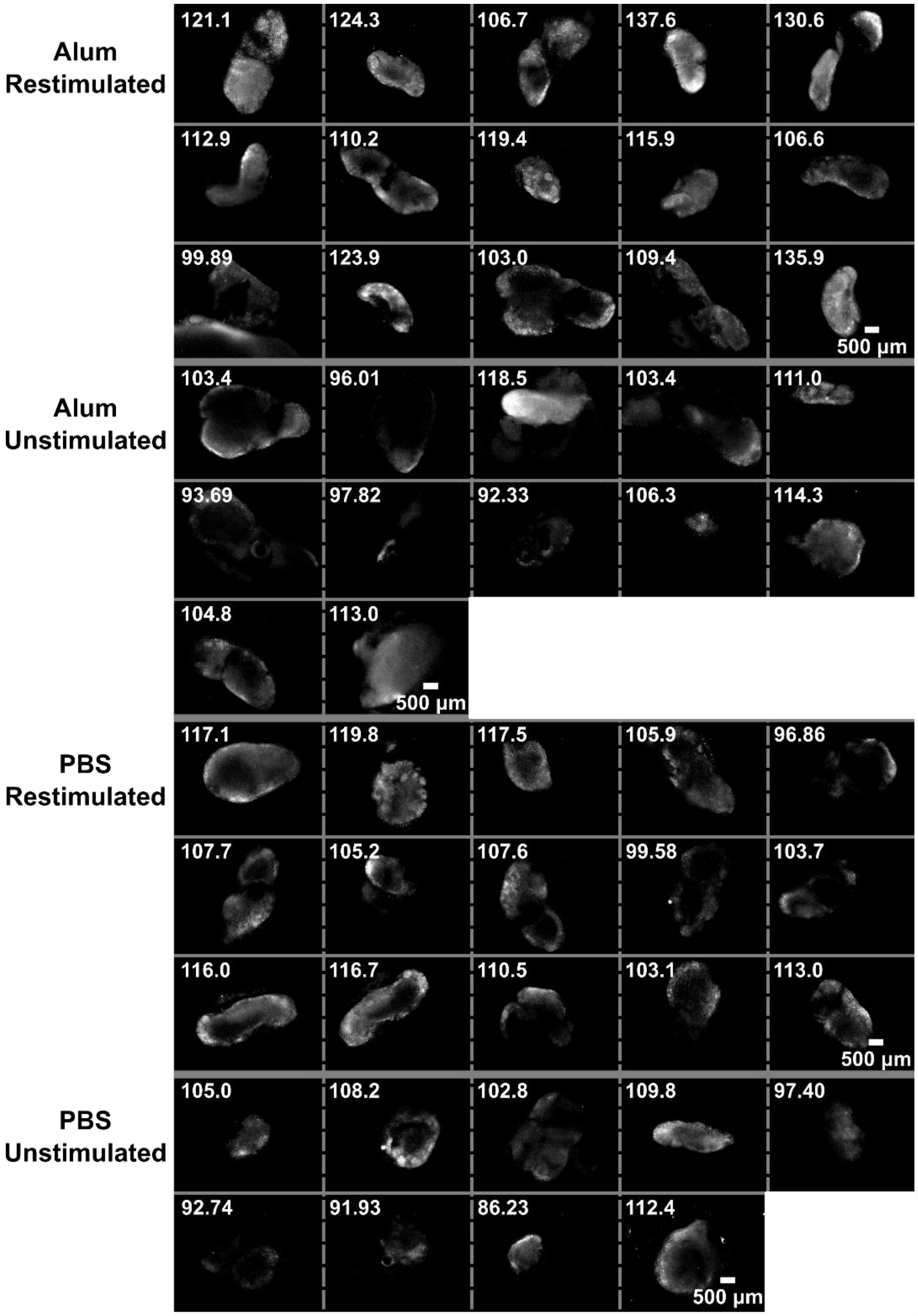


**Figure S9: Lymph node slices from vaccinated mice responded to protein-antigen challenge by upregulating surface expression of inflammatory markers.** Mice received i.v. OTII CD4+ T cells, then were immunized with either OVA+Alum (noted as Alum in the figure) or PBS. Slices from draining lymph nodes were collected on Day 4 and cultured ex vivo for 24 hours with either whole-protein OVA (Restimulated) or PBS (Unstimulated). Slices were then stained with AF647-CD69. Mean grey values for the CD69 channel is reported in each image.

**Table S1:** Gene targets from Qiagen inflammatory gene array. Relative Expression indicates the ratio of gene expression of slices/cell culture, where the gene expression in each condition was determined relative to the average expression of 5 housekeeper genes. Colors are matched to those in Figure 4: "Not Expressed" denotes genes not expressed in either sample (slices or cell culture). "1SD Different" and "2SD Different" indicate genes designated "Differentially Expressed" according to a threshold of either 1 or 2 std dev from the average Relative Expression (slices/cell culture), respectively. "Not Different" denotes genes expressed in both conditions and not statistically different under either threshold.

| Gene | Relative Expression (Sliced/Crushed) | Differential Designation | Gene | Relative Expression (Sliced/Crushed) | Differential Designation |
|---|---|---|---|---|---|
| Aimp1 | 8.94 | Not Different | Il10ra | 4.95 | Not Different |
| Bmp2 | 0.01 | 2SD Different | Il10rb | 26.37 | 1SD Different |
| Ccl1 | 0.32 | Not Expressed | Il11 | 33.37 | 2SD Different |
| Ccl11 | 10.33 | Not Different | Il13 | 17.25 | Not Different |
| Ccl12 | 22.80 | 1SD Different | Il15 | 0.21 | Not Different |
| Ccl17 | 23.80 | 1SD Different | Il16 | 0.02 | Not Expressed |
| Ccl19 | 3.07 | Not Different | Il17a | 0.02 | Not Expressed |
| Ccl2 | 385.30 | 2SD Different | Il17b | 0.00022 | 2SD Different |
| Ccl20 | 5.21 | Not Different | Il17f | 0.02 | Not Expressed |
| Ccl22 | 13.63 | Not Different | Il1a | 0.26 | Not Different |
| Ccl24 | 1.19 | Not Different | Il1b | 17.40 | Not Different |
| Ccl3 | 1.88 | Not Different | Il1r1 | 0.61 | Not Different |
| Ccl4 | 108.01 | 2SD Different | Il1rn | 150.13 | 2SD Different |
| Ccl5 | 1.70 | Not Different | Il21 | 0.02 | Not Expressed |
| Ccl6 | 0.60 | Not Expressed | Il27 | 0.02 | Not Expressed |
| Ccl7 | 4.42 | Not Different | Il2rb | 0.02 | Not Expressed |
| Ccl8 | 490.67 | 2SD Different | Il2rg | 0.29 | Not Expressed |
| Ccl9 | 12.06 | Not Different | Il3 | 0.02 | Not Expressed |
| Ccr1 | 1.80 | Not Different | Il33 | 3.04 | Not Different |
| Ccr10 | 0.02 | Not Expressed | Il4 | 3.29 | Not Different |
| Ccr2 | 0.02 | Not Expressed | Il5 | 4.66 | Not Different |
| Ccr3 | 0.77 | Not Expressed | Il5ra | 0.02 | Not Expressed |
| Ccr4 | 0.02 | Not Expressed | Il6ra | 8.73 | Not Different |
| Ccr5 | 5.96 | Not Different | Il6st | 33.41 | 2SD Different |
| Ccr6 | 0.02 | Not Expressed | Il7 | 1.38 | Not Different |
| Ccr8 | 0.02 | Not Expressed | Lta | 0.04 | Not Expressed |
| Cd40lg | 8.90 | Not Different | Ltb | 18.25 | 1SD Different |
| Csf1 | 0.27 | Not Different | Mif | 8.66 | Not Different |
| Csf2 | 8.28 | Not Different | Nampt | 0.56 | Not Expressed |
| Csf3 | 22.69 | 1SD Different | Osm | 1.55 | Not Different |
| Cx3cl1 | 0.02 | Not Expressed | Pf4 | 1.76 | Not Different |
| Cxcl1 | 0.27 | Not Expressed | Spp1 | 33.49 | 2SD Different |
| Cxcl10 | 803.88 | 2SD Different | Tnf | 0.02 | Not Expressed |
| Cxcl11 | 0.023 | Not Different | Tnfrsf11b | 0.02 | Not Expressed |
| Cxcl12 | 0.61 | Not Different | Tnfsf10 | 0.13 | Not Different |
| Cxcl13 | 56.92 | 2SD Different | Tnfsf11 | 0.02 | Not Expressed |
| Cxcl15 | 0.80 | Not Different | Tnfsf13 | 0.02 | Not Expressed |
| Cxcl5 | 3.03 | Not Different | Tnfsf13b | 0.01 | 2SD Different |
| Cxcl9 | 27.69 | 1SD Different | Tnfsf4 | 1.37 | Not Different |
| Cxcr2 | 0.02 | Not Expressed | Vegfa | 0.06 | Not Different |
| Cxcr3 | 0.02 | Not Expressed | Actb | | Housekeeper |
| Cxcr5 | 11.05 | Not Different | B2m | | Housekeeper |
| Fasl | 1.58 | Not Different | Gapdh | | Housekeeper |
| Ifng | 1.10 | Not Different | Gusb | | Housekeeper |
| | | | Hsp90ab1 | | Housekeeper |

**Table S2:** Fluorescent antibodies used to label cells for both flow cytometry and fluorescence microscopy

| TARGET | CLONE | FLUOROPHORE | PRODUCT NUMBER | VENDOR |
|---|---|---|---|---|
| ***B220** | RA3-6B2 | Pacific Blue | 103230 | Biolegend |
| ***B220** | RA3-6B2 | FITC | 103205 | Biolegend |
| ***B220** | RA3-6B2 | eFluor 570 | 41-0452-80 | eBioscience |
| **CD3** | 17A2 | Brilliant Violet 421 | 100227 | Biolegend |
| **CD3** | 17A2 | Brilliant Violet 510 | 100233 | Biolegend |
| **CD80** | 16-10A1 | Alexa Fluor 488 | 104715 | Biolegend |
| **CD11C** | N418 | PE | 117307 | Biolegend |
| **CD69** | H1.2F3 | PE-Cy7 | 104511 | Biolegend |
| **CD69** | H1.2F3 | PE | 104507 | Biolegend |
| ***CD69** | H1.2F3 | Alexa Fluor 647 | 104517 | Biolegend |
| **CD4** | GK1.5 | APC-Cy7 | 100525 | Biolegend |
| ***CD4** | GK1.5 | FITC | 100405 | Biolegend |
| **CD8** | 53-6.7 | Alexa Fluor 488 | 100726 | Biolegend |
| **CD25** | PC61 | Brilliant Violet 421 | 102033 | Biolegend |
| ***CD45** | 30-F11 | Alexa Fluor 488 | 103121 | Biolegend |
| ***LYVE-1** | ALY7 | eFluor 660 | 50-0443-82 | eBioscience |
| ***LYVE-1** | ALY7 | eFluor 570 | 41-0443-82 | eBioscience |
| ***F4/80** | BM8 | Brilliant Violet 605 | 123133 | Biolegend |
| ***CD169** | 3D6.112 | Alexa Fluor 594 | 142416 | Biolegend |
| ***KLH (Rat IgG 2a, k)** | RTK2758 | Alexa Fluor 488 | 400525 | Biolegend |

*Used for fluorescent imaging
